# Gene expression noise dynamics unveil functional heterogeneity of ageing hematopoietic stem cells

**DOI:** 10.1101/2022.08.04.502776

**Authors:** Reyna Edith Rosales-Alvarez, Jasmin Rettkowski, Josip Stefan Herman, Gabrijela Dumbović, Nina Cabezas-Wallscheid, Dominic Grün

## Abstract

Variability of gene expression due to stochasticity of transcription or variation of extrinsic signals, termed biological noise, is a potential driving force of cellular differentiation. While unicellular organisms exploit noise as a bet-hedging strategy, its role during multilineage differentiation of stem cells is underexplored. Utilizing single-cell RNA-sequencing to reconstruct cell state manifolds, we developed VarID2, a method for the quantification of biological noise at single-cell resolution. VarID2 reveals enhanced nuclear versus cytoplasmic noise across cell types of the peripheral blood, and distinct regulatory modes stratified by correlation between noise, expression, and chromatin accessibility. Noise levels are minimal in murine hematopoietic stem cells and increase during both differentiation and ageing. Differential noise identified myeloid-biased Dlk1+ long-term-HSCs in aged mice with enhanced quiescence and self-renewal capacity. VarID2 reveals fundamental properties of noise across cellular compartments, during stem cell differentiation and ageing, and uncovers distinct cellular sub-states invisible to conventional gene expression analysis.

## Introduction

Single-cell genomics has become a powerful method of choice for the identification of cell types and for the inference of tissue composition at single-cell resolution (Grün and van Oudenaarden, 2015; Kharchenko, 2021; Stegle et al., 2015). The reconstruction of cell state manifolds paired with pseudotime analysis facilitates the derivation of ancestral relations between cell states and enables the prediction of cellular differentiation trajectories (Sagar and Grün, 2020). However, such inference methods heavily rely on transcriptome similarity, and are therefore limited in their ability to capture control mechanisms of cell fate choice driven by lowly expressed genes. For example, single-cell lineage tracing by random cellular barcoding revealed early lineage priming of hematopoietic stem cells (HSCs) (Weinreb et al., 2020), which remained undetected when relying on single-cell RNA-seq (scRNA-seq) data alone. A known problem for the quantification of subtle expression changes, in particular for lowly expressed genes, is the substantial level of technical variability masking genuine biological variability, or noise (Brennecke et al., 2013; Grün et al., 2014).

Gene expression noise is prevalent in unicellular organisms (Elowitz et al., 2002; Ozbudak et al., 2002) and can underlie bi-stable systems such as the *E. coli* lac operon (Ozbudak et al., 2004). Increased variability of gene expression has been observed during *in vitro* differentiation of embryonic stem cells (Stumpf et al., 2017) or upon reprogramming of induced pluripotent stem cells (Buganim et al., 2012), yet its role during cell fate decision within multilineage systems *in vivo* is underexplored (Eling et al., 2019; Raj and van Oudenaarden, 2008).

Although scRNA-seq allows noise quantification within homogenous cell populations (Grün et al., 2014; Kar et al., 2017; Kim et al., 2015; Kolodziejczyk et al., 2015; Vallejos et al., 2015), available methods cannot capture biological noise dynamics at high resolution across complex cell state manifolds.

We recently proposed VarID as a method for quantifying local gene expression variability in cell state space, which eliminates the mean dependence of gene expression variability but does not explicitly distinguish technical and biological sources of noise (Grün, 2020). We here introduce VarID2 to overcome this major limitation by modeling defined sources of technical noise in local cell state neighborhoods, facilitating the inference of actual biological variability. We demonstrate that VarID2 predicts biological noise levels consistent with state-of-the-art Bayesian noise models (Eling et al., 2018; Vallejos et al., 2015), which are only applicable to pairwise comparisons of large homogenous cell populations profiled by scRNA-seq, and, hence, do not permit the investigation of noise dynamics during cellular differentiation.

VarID2 analysis of human peripheral blood mononuclear cells (PBMCs) indicates a general increase of transcriptome variability in the nucleus compared to the cytoplasm, and the relation between chromatin accessibility, gene expression, and noise uncovers distinct modes of gene regulation.

Noise quantification within the murine hematopoietic system reveals minimal noise in HSCs which increases upon differentiation. The hematopoietic system is known to be affected by ageing, with a gradual functional decline of HSCs and an emerging myeloid bias in the bone marrow (De Haan and Lazare, 2018). It is still unclear to what extent the age-dependent decline of the hematopoietic system can be attributed to an emerging HSC heterogeneity, resulting from age-related changes of cell-intrinsic properties, or from a changing bone marrow microenvironment. By applying VarID2 we observed increased transcriptome variability in HSCs of aged mice. The top noisy gene in aged HSCs, *Dlk1*, facilitates the discrimination of two sub-populations of HSCs which are almost indistinguishable on the global transcriptome level, yet exhibit clear differences in terms of quiescence, self-renewal capacity, and myeloid bias. We argue that age-related emergence of Dlk1+ HSCs with cell-intrinsic myeloid bias could contribute to the age-dependent change of bone marrow composition. Hence, we demonstrate that single-cell resolution analysis of gene expression noise can yield fundamentally new biological insights. VarID2 was integrated into our RaceID toolkit for single-cell analysis publicly available on CRAN.

## Results

### Modeling local gene expression variability in cell state space

To model local gene expression variability in cell state space, we are building upon our previous VarID method (Grün, 2020). VarID constructs a pruned k-nearest neighbor (knn) graph in cell state space and tests transcript count differences between the “central” cell and each of its neighbors against a background model. In VarID2, this background model is defined as a negative binomial with a local mean of raw unique molecular identifier (UMI) counts and a corresponding standard deviation obtained from a local fit of the mean-variance dependence across all genes (Methods). To overcome the lack of VarID in resolving technical and biological noise components and to estimate genuine biological variability, we reasoned that two sources of noise dominate the observed UMI count variance measured for each gene in such local neighborhoods. At low expression, sampling noise, i.e., binomial variance captures the average trend (Figure 1A): in this regime, the dependence of the coefficient of variation (CV) on the mean follows a line of slope −1/2 in logarithmic space. At high expression, the CV-mean dependence saturates and approaches a baseline variance level. As described previously (Grün et al., 2014), this baseline is determined by the shared variability affecting all genes equally, which we refer to as total UMI count variability. Major sources of this noise component are cell-to-cell differences in sequencing efficiency and cell volume. We inferred this noise component by fitting a Gamma distribution to the total UMI count distribution across cells in each local neighborhood. The resulting Poisson-Gamma mixture corresponds to a negative binomial distribution, describing the UMI counts *X_i,j_* for each gene *i* across a neighborhood with central cell *j*. The parameters of this distribution are the local mean *μ_i,j_* of the raw UMI counts, and dispersion parameter *r^t^_j_* given by the rate parameter of the Gamma distribution (Methods).

**Figure 1.**
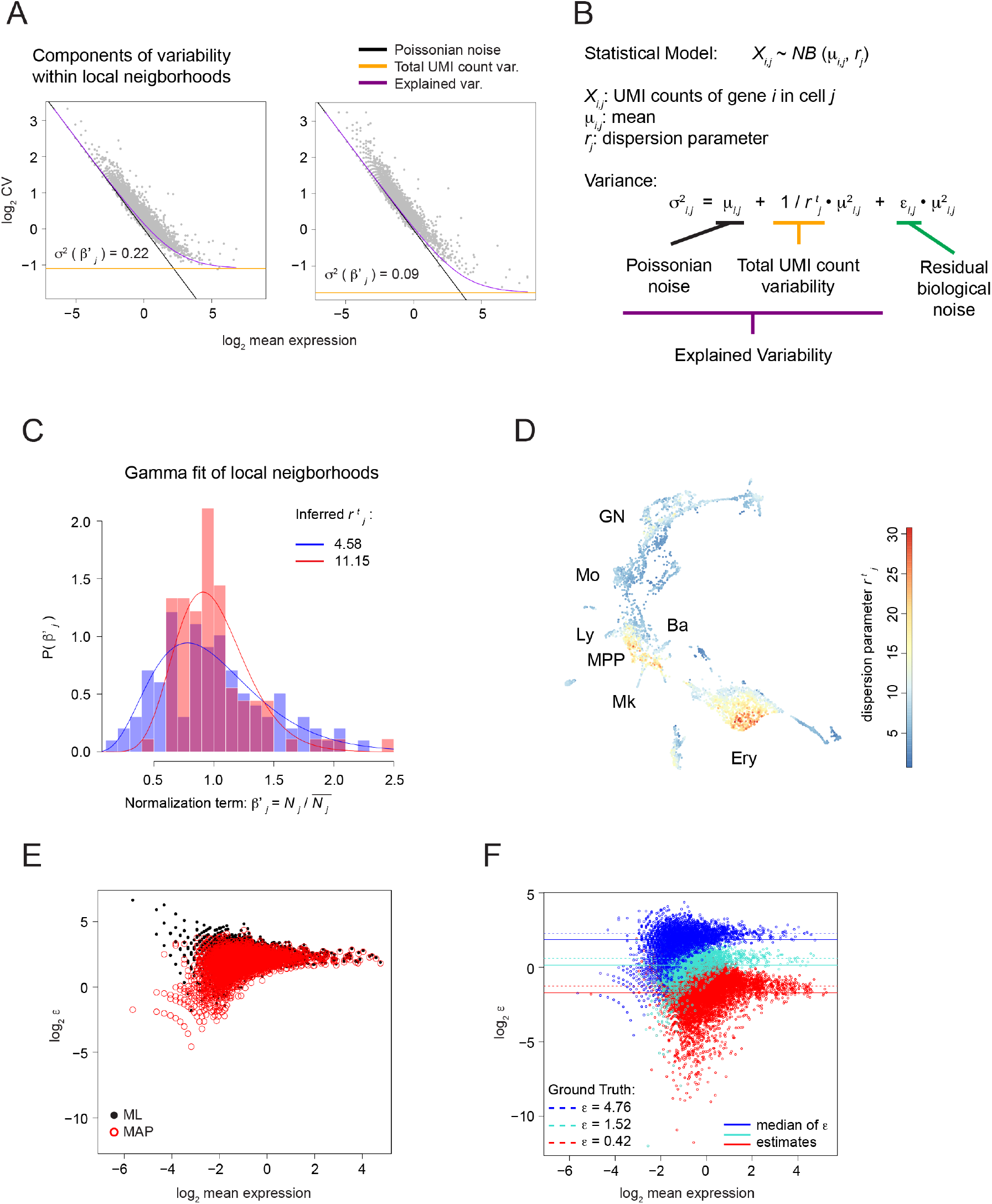
Local decomposition of gene expression noise in cell state space. (A) Coefficient of variation as a function of the mean expression on logarithmic scale. The explained variability and its components, Poissonian noise and total UMI count variability, are highlighted. Plots correspond to two individual neighborhoods of 101 cells each from a Kit+ hematopoietic progenitor dataset (Tusi et al., 2018). (B) Negative Binomial model for the UMI counts *X_i,j_*. The variance is split into three components: Poissonian noise, total UMI count variability, and residual biological noise. (C) Estimation of the dispersion parameter 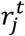 for two individual neighborhoods shown in (A). Mean-normalized total UMI counts *β′_j_* are fitted by a gamma distribution, with shape parameter 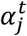 equal to the dispersion parameter 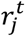 in (B). (D) UMAP plot highlighting 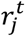 estimates across the hematopoietic progenitor dataset. MPP, multipotent progenitors; Ly, lymphocytic; Mo, monocytic; GN, granulocytic neutrophil; Ba, basophylic; Mk, megakaryocytic; Ery, erythroid. (E) Comparison of *ε* estimates obtained by Maximum Likelihood (ML) estimation (black) and Maximum a posteriori (MAP) estimation (red). A simulated dataset with three levels of gene expression noise was used (see Methods and Figure S1B). Here, only *ε* estimates corresponding to the highest noise level are shown. (F) *ε* estimates for a simulated dataset with three different biological noise levels (Methods). Colours highlight group of genes with different simulated biological noise levels (low, medium, or high). Simulated ground truths of noise values (dashed lines), and median values of the *ε* estimates (solid lines) are indicated for each group. Hyperparameter *γ* = 1.

The remaining residual variability in excess of these two major sources can be summarized into an additional dispersion parameter *ε_i,j_* (Figure 1B). We refer to this residual variability as biological noise since it captures gene-specific deviations from the global trend determined by sampling variability and total UMI count variance.

By applying VarID2 to scRNA-seq data of mouse Kit+ hematopoietic progenitors (Tusi et al., 2018) comprising major branches of erythrocyte and neutrophil progenitors, we observed that *r^t^j* indeed varies substantially between distant neighborhoods in cell state space. Thus, a local noise model is required to quantify this noise component for heterogeneous cell populations (Figure 1C and 1D). Since a maximum-likelihood (ML) fit of the biological noise *ε_i,j_* led to inflated estimates for lowly expressed genes, we incorporated a weakly informative Cauchy prior and performed maximum a posterior (MAP) estimation of *ε_i,j_*, which eliminated the inflation (Methods and Figure 1E and S1A). To test our noise model quantitatively, we simulated cell neighborhoods with defined technical and biological noise levels based on gene expression parameters from (Tusi et al., 2018) (Methods and Figure S1B). We optimized the scale parameter *γ* of the Cauchy prior by jointly matching the median and minimizing the standard deviation of the estimates compared to the simulated ground truth (Figure S1C). This analysis demonstrates the accuracy of our noise estimates across three different noise levels, as well as the absence of a systematic mean-variance dependence (Figure 1F).

In order to make VarID2 scalable we restricted the model to MAP estimation of the residual noise parameter. BASiCS has been introduced as a full Bayesian noise model with multiple parameters (Eling et al., 2018; Vallejos et al., 2015), yet this model is computationally expensive and application to a larger number of local neighborhoods is infeasible. Moreover, the biological noise parameter of BASiCS (Eling et al., 2018) is defined as the residual over-dispersion from the average mean-variance dependence. In contrast, VarID2 assigns a clear interpretation to *ε_i,j_* as a residual after deconvoluting defined noise components. Reassuringly, *ε_i,j_* is highly correlated with BASiCS’ over-dispersion parameter (Pearson’s correlation coefficient 0.85) with diminished correlation of the ML estimate (Pearson’s correlation coefficient 0.79), supporting our choice of the prior (Figure S1D and S1E). However, although BASiCS correctly discriminates different noise levels, the estimates deviate from the simulated parameters (Figure S1F). Hence, VarID2 overcomes limitations of available methods for the noise quantification across large numbers of local neighborhoods, enabling the analysis of noise differences between multiple populations and along differentiation trajectories.

### Nuclear versus cellular transcripts exhibit elevated noise levels in peripheral blood mononuclear cells

We first applied VarID2 to test the hypothesis that nuclear export of mRNAs serves as a buffer to reduce transcriptional noise, as described for a limited set of genes measured by single-molecule fluorescent in situ hybridization (smFISH) in HeLa cells and primary Keratinocytes (Battich et al., 2015). To compare transcriptional noise of nuclear and cytoplasmic transcripts on a genome-wide scale across a number of different cell types, we ran VarID2 on scRNA-seq and single-nucleus RNA-seq (snRNA-seq) data of human peripheral blood mononuclear cells (PBMCs, datasets generated by 10x Genomics, see Table S1). For both datasets VarID2 identified monocytes, NK cells, T cells, and B cells, which could be further sub-classified into different sub-types consistently observed in both datasets (Figure 2A, 2B and S2A-C). Across all cell populations, naïve T cells were found to exhibit minimal noise levels suggesting that transcriptional variability is reduced in less differentiated cell states (Figure 2C and 2D). To enable the comparison of cell populations between the two datasets, we annotated corresponding cell types based on data integration with Harmony and the Seurat pipeline (Methods and Figure S2D). Substantial noise reduction in cellular versus nuclear transcripts was consistently observed across all cell types and for the majority of all genes (Figure 2E and S2E) independently of the expression level (Figure 2F and S2F), indicating that nuclear export could indeed facilitate noise reduction on a genome-wide level across cell types.

**Figure 2.**
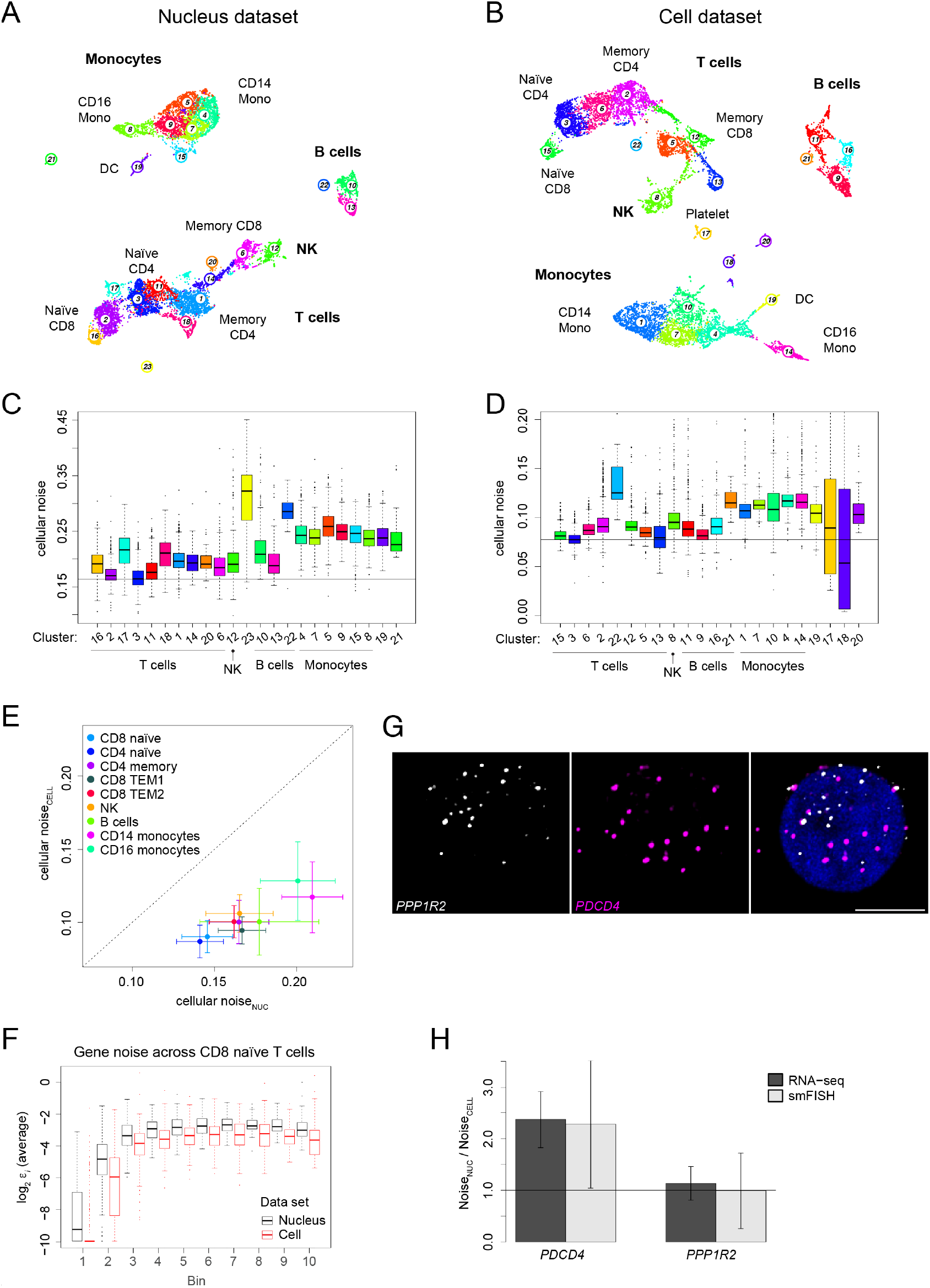
Elevated noise levels of nuclear versus whole-cell transcriptomes in human PBMCs. (A) Clustering and UMAP representation of single-nuclei RNA-seq data, consisting of human peripheral blood mononuclear cells (PBMC) profiled with the Single Cell Multiome kit from 10x Genomics (See Table S1). (B) Clustering and UMAP representation of single-cell RNA-seq data, comprising human PBMCs (10x Genomics). (C) Quantification of cellular noise (average *ε* across all genes per cell) across clusters shown in (A). The horizontal line corresponds to the median of CD4 naïve T cell estimates (cluster 3), exhibiting reduced noise levels. Boxes indicate inter-quartile range (IQR), and whiskers correspond to ±1.5*IQR of the box limits. Outliers beyond the whisker limits are depicted. Vertical axis limits are manually adjusted for better visualization. (D) Same as (C), but for cellular noise estimates for the single-cell dataset (see (B)). The horizontal line corresponds to the median of CD4 naïve T cell estimates (cluster 3). (E) Comparison of cellular noise levels between both datasets. The dot plot shows the average cellular noise per cluster and their corresponding standard deviation (error bars). Horizontal axis corresponds to the estimates of the nucleus data and the vertical axis to the cell data estimates. Selection of similar cell populations between both datasets was performed by dataset integration using Harmony (Korsunsky et al., 2019). See Figure S3D. (F) Comparison of *ε* estimates per gene across CD8 naïve T cells between nucleus and cell datasets. Genes that do not change expression across compartments were selected and grouped into ten equally populated bins, based on their mean expression. See also Figure S3F. Boxes indicate interquartile range (IQR), and whiskers correspond to ±1.5*IQR of the box limits. Outliers beyond the whisker limits are depicted. (G) Expression of *PDCD4* (elevated nuclear noise) and *PPP1R2* (similar noise levels in nucleus and whole cell) was quantified by smFISH in human CD8 naïve T cells (see also Figure S2H). Representative images of maximum intensity projections are shown. DAPI in blue, scale bar is 5 μm. (H) Comparison of VarID2 noise estimates to smFISH-derived values. The noise ratio between nuclear and cellular compartments is shown. Error bars correspond to standard error (Methods). DC, dendritic cells; NK, natural killer cells; TEM, effector memory T cells; Mono, monocytes.

To validate the observation of increased noise of nuclear versus cellular transcripts, we quantified mRNA abundance by smFISH on CD8 naïve T cells isolated from human peripheral blood. We selected candidate genes with equal expression in the nuclear and the cellular compartment (Figure S2F). The translational inhibitor programmed cell death-4 (*PDCD4*), involved in cell apoptosis and also in the control of CD8 T cell activation (Hilliard et al., 2006) exhibits increased noise in the nucleus according to our prediction (Figure S2G). Moreover, *PDCD4* undergoes alternative splicing and one of its isoforms is regulated by nuclear retention (Park and Jeong, 2016). This suggests that post-transcriptional regulatory mechanisms may mediate elevated nuclear noise. In contrast, the gene encoding phosphatase inhibitor 2 (*PPP1R2*) was predicted to exhibit similar nuclear and cellular noise levels (Figure S2G). For these genes, we quantified nuclear and cytoplasmic mRNA counts by smFISH (Figure 2G and S2H), and computed the ratio of residual biological noise between nucleus and whole cells, which was in excellent agreement with the noise ratios predicted by VarID2 (Methods and Figure 2H).

### Co-analysis of chromatin accessibility and gene expression noise reveals distinct modes of gene regulation

In order to gain insights into the influence of chromatin accessibility on gene expression noise, we analyzed a multiomics PBMC dataset (see Table S1), which combines snRNA-seq and single-cell Assay for Transposase-Accessible Chromatin sequencing (scATAC-seq) from the same cell, by using the Signac package (Stuart et al., 2021). We focused our analysis on the chromatin accessibility across individual genes at two levels, gene activity and individual peak signal.

Gene activity was defined as the sum of detected fragments across all peaks located in the gene body and 2kb upstream of the transcriptional start site (TSS). We computed Pearson correlations for paired comparisons of expression (Ex), noise (N), and gene activity (GA) (Figure S3A). In agreement with the general notion that open chromatin promotes gene expression, we observed a substantial number of genes with a positive correlation between expression and gene activity (1,623). However, the majority of the genes did not exhibit a clear association, potentially due to the sparsity of at least one of the modalities.

Similarly, a substantial number of genes showed a positive correlation between gene activity and noise (933, Figure S3A), and the majority of those also displayed a positive correlation of expression and gene activity (857) (termed class A genes, Figure 3A). On the other hand, most genes with a negative correlation between noise and gene activity (95) exhibited a positive expression – gene activity correlation (84) (termed class B genes, Figure 3A).

**Figure 3.**
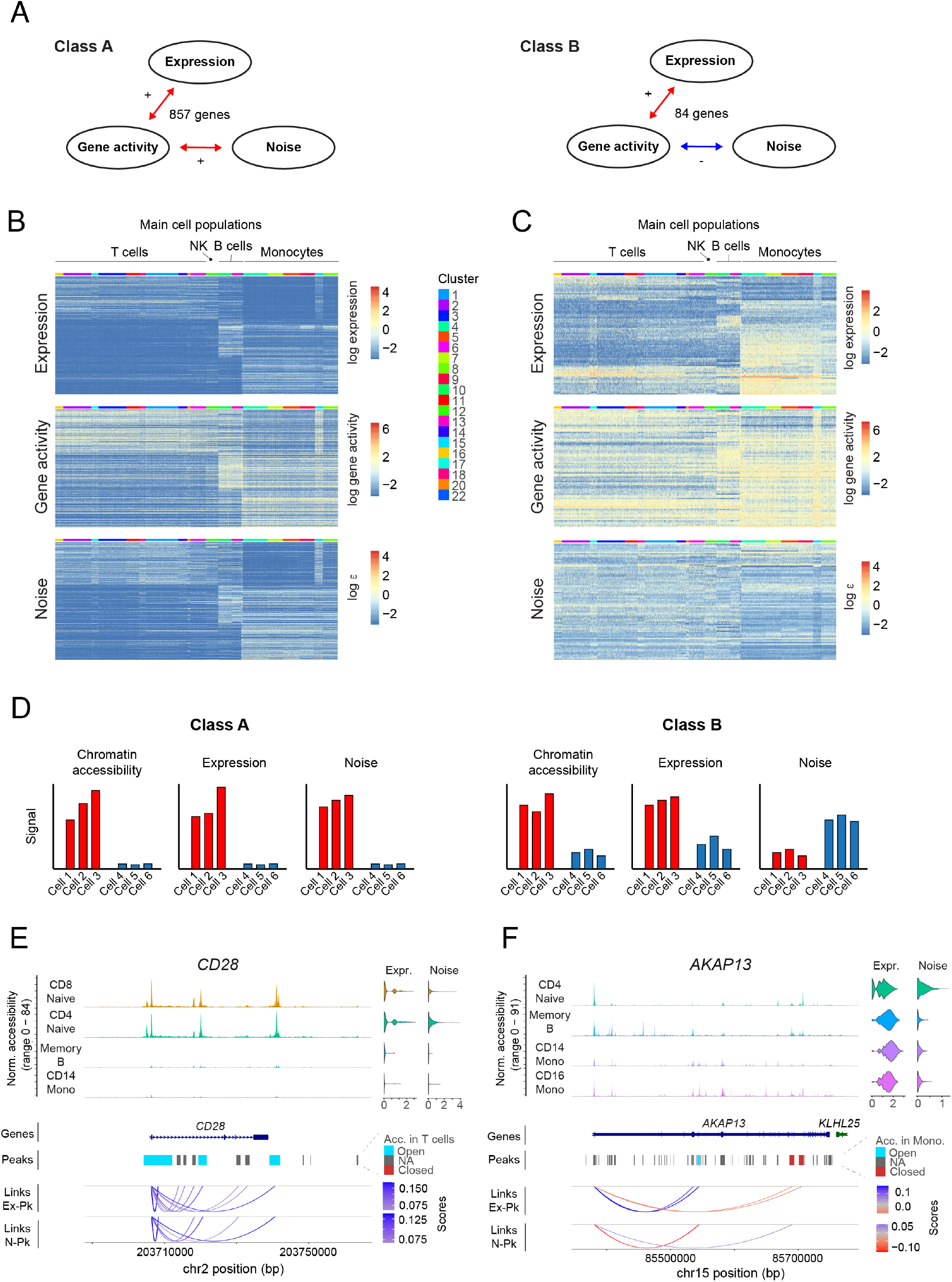
Joint analysis of chromatin accessibility and gene expression noise reveals gene modules with distinct noise regulation. (A) Two sets of genes were analyzed based on the correlations in Figure S3A. Class A genes (left side) have positive expression – gene activity and noise – gene activity correlations. Class B genes have positive expression – gene activity correlation but negative noise – gene activity correlation. (B) Patterns of expression (top), gene activity (middle) and noise (down) of genes belonging to class A. For convenience only ~300 genes are shown. See also Figure S3B for heatmaps with the whole set of these genes. (C) Similar as (B), but showing genes of class B. All genes in this category were included. (D) Diagram summarizing the observed patterns in chromatin accessibility, expression and noise for the set of genes in class A and class B. See main text for further details. (E) Genomic region of *CD28* (class A gene). Upper panel: normalized accessibility signal, aggregated across cells from selected clusters. Violin plots (top right) show expression and noise levels across each cluster. Differential accessibility test of T cells against the remaining dataset was performed. Peaks (middle panel) were annotated based on increased accessibility (“Open”), no change (“NA”) or decreased accessibility (“Closed”). Threshold values: logFC > 1.25, padj < 0.001. Gene linkages (Ma et al., 2020; Stuart et al., 2021) between expression and accessibility within individual peaks (links Ex-Pk) or noise and peak accessibility (links N-Pk) are shown in the lower panel, with scores corresponding to Pearson correlation coefficients. These links bind the TSS of the corresponding gene and peaks where a significant correlation was detected, and they do not represent spatial chromatin organization. (F) Similar as (E), but showing data of *AKAP13* (class B gene). Differential accessibility test was performed by comparing monocytes cells against the remaining dataset.

Class A genes tend to be expressed exclusively in either T cells, B cells, or monocytes, with high accessibility and noise signal in these cell types (Figure 3B and S3B), while class B genes exhibit a mixture of expression patterns. While most of these genes are dominantly expressed in a specific cell population, they are still expressed at lower levels in other cell types (Figure 3C). Other genes of class B are more ubiquitously expressed across the entire dataset. As expected, noise of class B genes is generally anti-correlated with expression. For the remaining genes (class C genes), noise and gene activity did not correlate (Figure S3C).

Hence, expression level and noise increase with chromatin accessibility for class A genes, suggesting that these genes exhibit an on-off pattern without precise control of the transcriptional level (Figure 3D). In contrast, genes in class B show reduced variability when chromatin becomes more accessible and expression increases, and may thus require more precise regulation of their transcriptional output (Figure 3D).

Pathway enrichment analysis for these sets of genes allows to assess whether they are involved in particular cellular functions. For each main cell population (T cells, B cells and monocytes), we performed enrichment analysis over a complete list of marker genes obtained by differential gene expression analysis (Methods), selecting subsets of marker genes found in class A, or those that do not belong to class A (Figure S3D). For the three cell types, marker genes belonging to class A are significantly more enriched in cell type-specific immune signaling functions compared to the full list of marker genes. Among these enriched pathways we found, e.g., co-stimulation by the CD28 family for T cells, signaling by the B cell receptor for B cells, and interleukin 10 signaling for monocytes. In contrast, marker genes that do not belong to class A, yielded more general categories in case of T cells and monocytes, and no enrichment for B cells (Figure S3D). On the other hand, enrichment analysis for marker genes within class B did not return any pathway, suggesting that only particular genes within broader functional categories require precise control of transcriptional activity.

Furthermore, we investigated associations reflected by correlations between expression or noise, respectively, and fragments at the level of individual peaks by following a recently proposed strategy (Ma et al., 2020) implemented in Signac (Stuart et al., 2021). This method addresses confounding factors such as GC content and sequence length by comparing the peak-gene correlation against a background signal and testing the significance of the correlation. We adapted the input in order to obtain both, expression – peak signal (Ex – Pk) and noise – peak signal (N – Pk) correlations. For simplicity, we focused our analysis on peaks falling around the TSS and gene body, setting aside potential regulatory regions in *cis*.

Focusing on class A and B genes, we observed consistent patterns at the level of specific peaks as compared with gene activity signal. Genes of class A show an enrichment in both positive expression - peak and noise - peak correlations with substantial overlap of these peaks.

For instance, the T cell co-receptor *CD28* (belonging to class A) exhibits common links of positive expression – peak and noise – peak correlation (Figure 3E). A complementary behavior was observed for the class B gene *AKAP13* which shows peaks with positive expression – peak but negative noise – peak correlation, and vice versa (Figure 3F).

Moreover, peaks within the *CD28* locus exhibiting positive expression – peak and noise – peak correlations are differentially accessible in T cells versus other cells (Figure 3E). Likewise, peaks associated with increased expression and low noise across the *AKAP13* sequence exhibit increased accessibility in monocytes, while peaks associated with decreased expression and high noise are less accessible (Figure 3F). Therefore, correlations at the higher level of gene activities largely reflect the dynamics at individual peaks, supporting the distinction between class A and class B genes.

Taken together, gene expression noise can discern different modes of gene regulation corresponding to noisy on/off switches (class A) versus tight regulation of expression levels (class B).

### Gene expression noise increases during hematopoietic differentiation

To interrogate dynamics of gene expression noise during multilineage stem cell differentiation, we focused on the hematopoietic system and analyzed a dataset of ~44,000 mouse Kit+ hematopoietic progenitors, covering LT-HSCs, multipotent progenitors (MPPs), and fate-committed progenitors of all major blood lineages (Dahlin et al., 2018). Cluster-to-cluster transition probabilities (Grün, 2020) predicted by VarID2 recapitulate the architecture of the hematopoietic tree (Figure 4A and S4A). LT-HSCs identified as the *Slamf1*+ *Ly6a*+ *Kit*+ *Cd34*^low^ *Cd48*^low^ cluster 10 exhibit the lowest averaged noise level (mean noise of all genes in a local neighbourhood) among all clusters (Figure 4B and S4B). Hence, transcriptional noise is suppressed in LT-HSCs, indicating a stable, transcriptionally homogenous stem cell state. For all lineages, we observed an increase of transcriptional noise with differentiation progress (Figure 4B and S4B). We further analyzed cell-to-cell transcriptome correlation within local neighborhoods (Figure S4B), and found that LT-HSCs are among the clusters with the highest Spearman correlation of single-cell transcriptomes. Hence, transcriptional variability in LT-HSCs is correlated across genes, suggesting fluctuations of entire gene modules.

**Figure 4.**
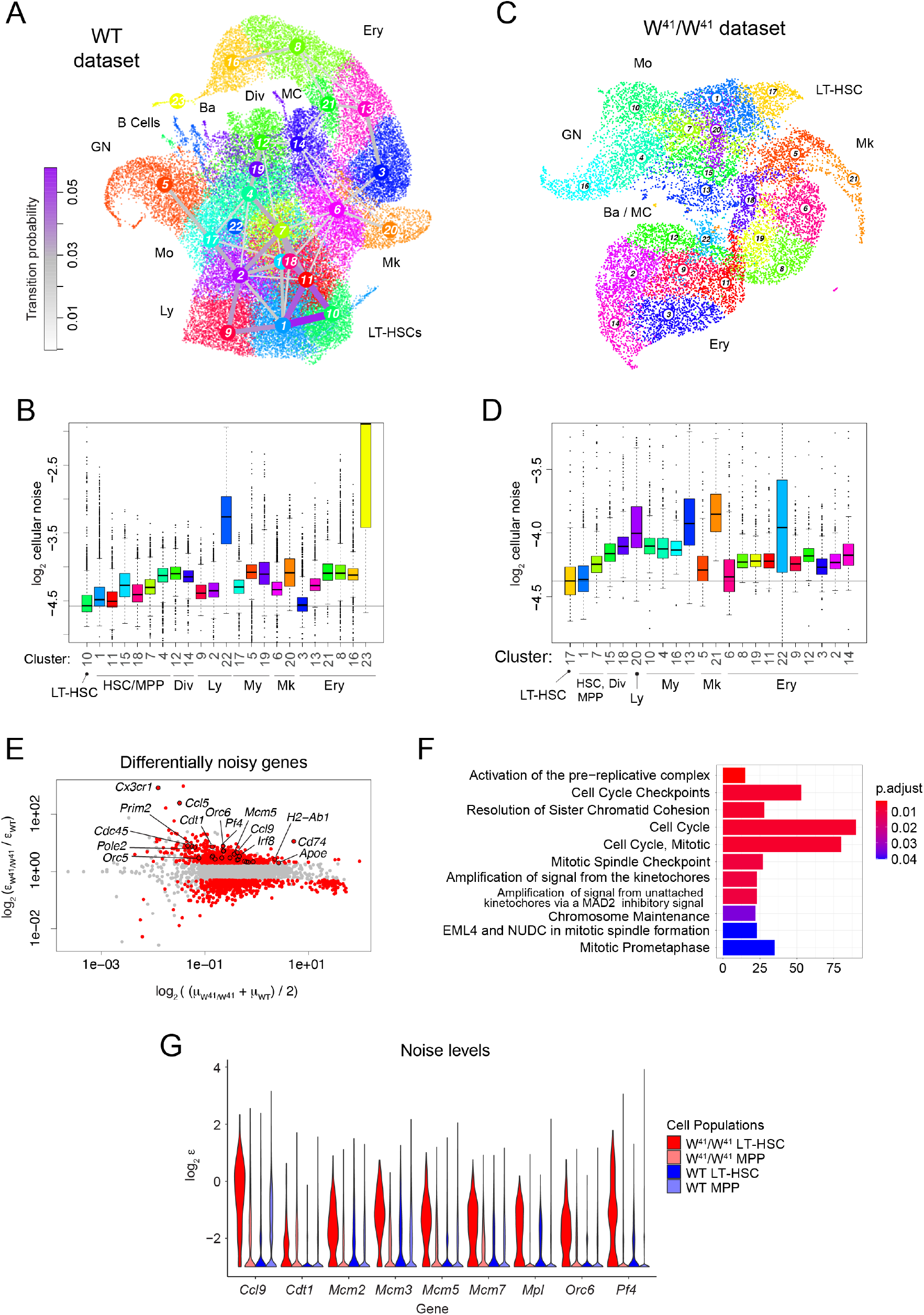
Gene expression noise increases during hematopoietic differentiation. (A) UMAP representation of hematopoietic stem and progenitor cells from the bone marrow of wildtype (WT) mice (Dahlin et al., 2018). Major cell populations and VarID2 transition probabilities (Methods) between clusters are highlighted. (B) Quantification of cellular noise (average *ε* across all genes per cell) across clusters from the WT dataset in (A). Horizontal line corresponds to the median noise level of the LT-HSC population. Boxes indicate inter-quartile range (IQR), and whiskers correspond to ±1.5*IQR of the box limits. Outliers beyond the whisker limits are depicted. Vertical axis limits are manually adjusted for better visualization. (C) UMAP representation of a hematopoietic stem and progenitor cells from *Kit* W^41^/W^41^ mutant mice (Dahlin et al., 2018). (D) Same as (B), but showing cellular noise estimates of the W^41^/W^41^ dataset in (C). (E) Differentially noisy genes identified between the LT-HCS populations of W^41^/W^41^ versus WT mice. MA plot shows log2FC of noise on the *y* axis, and average expression on the *x* axis. Threshold values: log2FC > 1, padj < 0.001. (F) Pathway enrichment analysis of the genes with increased noise in W^41^/W^41^ mice from (E). (G) Noise *ε* estimates of genes involved in DNA replication. Quantities from each dataset were separated into LT-HSCs and the remaining cells, denoted as MPP. LT-HSC, long-term hematopoietic stem cells; MPP, multipotent progenitors; Ly, lymphocytic; My, myelocytic; Mo, monocytic; GN, granulocytic neutrophil; Ba, basophylic; MC, mast cells; Mk, megakaryocytic; Ery, erythroid; Div; dividing cells.

Impaired Kit signaling affects long term repopulation capacity of HSCs (Sharma et al., 2007), and *in vitro* culture of W^41^/W^41^ mutant mice with impaired Kit kinase activity demonstrated reduced proliferation within the HSC compartment (Dahlin et al., 2018). To test whether stochastic activation of cell cycle genes could underlie the perturbed exit from quiescence, we performed VarID2 analysis of scRNA-seq data generated from W^41^/W^41^ mutant hematopoietic progenitors (Dahlin et al., 2018) (Figure 4C). We were able to identify all major hematopoietic lineages with perturbed relative abundances as reported in the original study. By matching cluster centers between wildtype and mutant datasets (Methods), we identified mutant cluster 17 as the unique match to the wildtype LT-HSC cluster 10 (Figure S4D), which was also supported by LT-HSC marker expression (Figure S4E). Similar to wildtype cells, mutant cells exhibit minimal noise levels in LT-HSCs and an increase upon differentiation (Figure 4D). We next interrogated noise differences between wildtype and mutant LT-HSCs based on differentially noisy genes (Figure 4E), and detected a strong enrichment of cell cycle genes (Figure 4F). In particular, several members of the pre-replication complex (*Mcm2, Mcm3, Mcm5, Mcm7, Orc6*) where among the top differentially noisy genes (Figure 4G) despite only small differences in expression levels (Figure S4F). These genes are required for the initiation of replication and showed elevated noise levels in LT-HSCs versus MPPs. Taken together, these observations suggest that cell cycle activation in W^41^/W^41^ mutant LT-HSCs becomes more stochastic. This is consistent with the observation of Dahlin et al., that the number of colonies obtained from *in vitro* culture is overall comparable between wildtype and W^41^/W^41^ mutants, yet the frequency of very small colonies was significantly increased, indicating the presence of LT-HSCs that fail to become fully proliferative. Hence, the noise analysis can generate hypotheses consistent with the observed perturbed proliferation phenotype in *Kit* mutant mice.

### Gene expression noise increases in LT-HSCs upon ageing

Ageing increases cell-to-cell variability of CD4+ T cells upon immune stimulation (Martinez-Jimenez et al., 2017). To test whether an increase of gene expression noise also occurs in HSCs upon ageing, and to investigate if this could explain observed phenotypic changes such as myeloid lineage bias (Geiger et al., 2013), we applied VarID2 to scRNA-seq data of HSCs isolated from young (2-3 months old) and aged (17-18 months old) mice (Hérault et al., 2021). In this study, sequencing was performed in two batches (denominated as A and B) of young and aged mice, which were separated by VarID2 clusters (Figure 5A and S5A). To avoid confounding of noise quantification by batch integration, we separately analyzed clusters corresponding to the two batches. We focused our analyses on the clusters maximizing expression of LT-HSC markers (*Hlf, Hoxa9, Mecom*) within each age group and batch (Figure S5B): cluster 7, 15, 1, and 6 for young A, young B, aged A, and aged B, respectively. Compared to multipotent progenitors (MPPs), these clusters show decreased noise (Figure 5B and S5C), consistent with the analysis of data from Dahlin et al.. For both batches, we observed elevated noise levels in aged versus young LT-HSCs (Figure 5C), indicating that the transcriptome of LT-HSCs becomes more variable with age.

**Figure 5.**
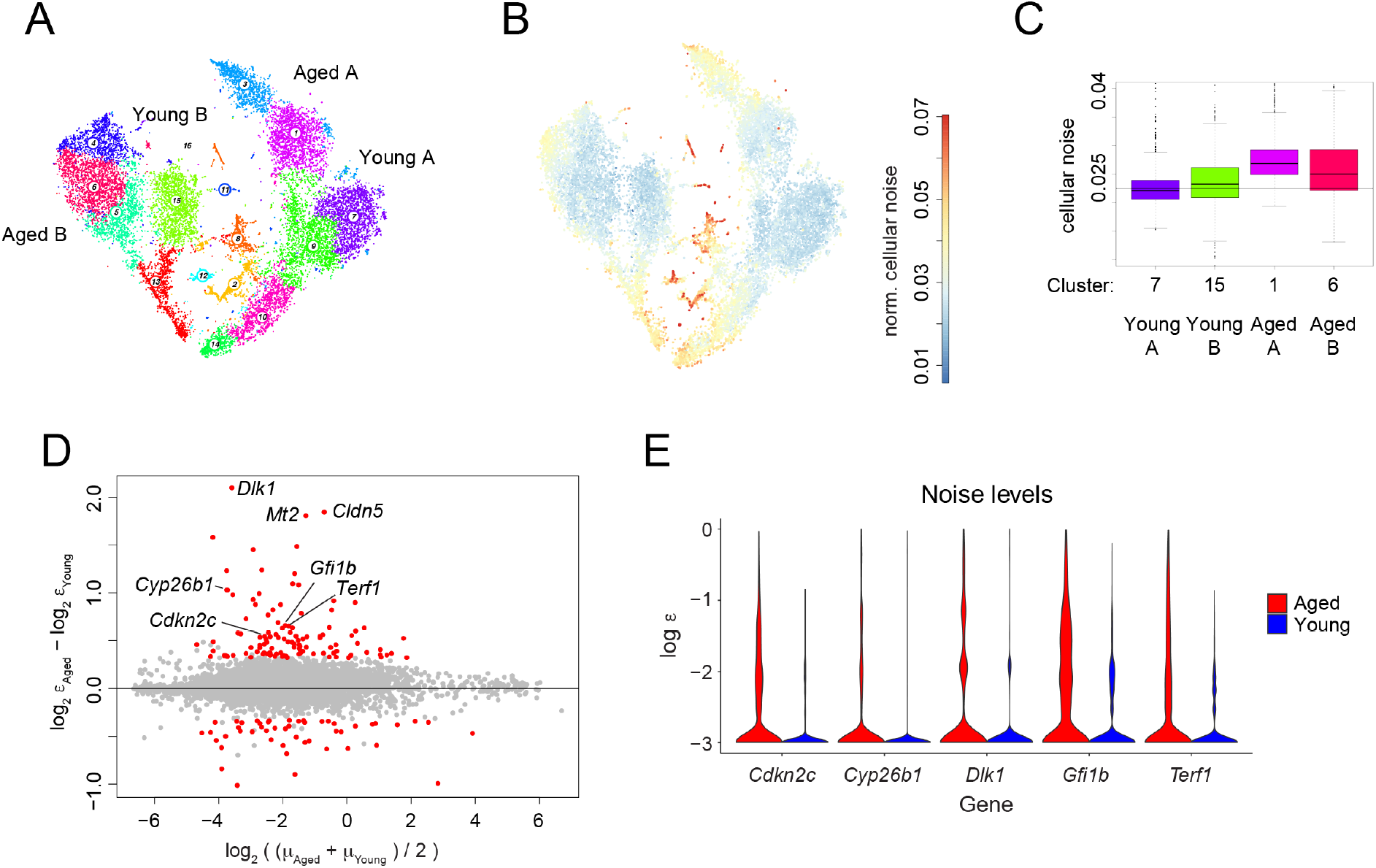
Gene expression noise increases in LT-HSCs upon ageing. (A) t-SNE representation of young and aged hematopoietic stem cells (Hérault et al., 2021), sequenced in two batches A and B (see also Figure S5A). LT-HSC populations identified based on marker gene expression for each condition and batch identity are highlighted (see also Figure S5B). (B) t-SNE plot highlighting cellular noise estimates across cells in the the Hérault et al. dataset. (C) Comparison of cellular noise across the four LT-HSC populations identified in (A). Boxes indicate inter-quartile range (IQR), and whiskers correspond to ±1.5*IQR of the box limits. Outliers beyond the whisker limits are depicted. Vertical axis limits are manually adjusted for better visualization. (D) Differentially noisy genes identified across LT-HCS populations, comparing aged versus young samples. MA plot shows log2FC of noise on the *y* axis, and average expression on the *x* axis. Threshold values: logFC > 1.25, padj < 0.001. (E) Noise *ε* estimates of some example genes detected as highly noisy in aged versus young LT-HSCs in (D).

Analysis of differentially noisy genes (Methods, Figure 5D) confirmed a larger number of genes with elevated noise in aged LT-HSCs. Among these genes we detected the inhibitor of Telomerase *Terf*, cell cycle suppressers such as cyklin-dependent kinase inhibitor *Cdkn2c*, and *Gfilb*, an essential regulator of erythro-megakaryopoiesis (van der Meer et al., 2010) (Figure 5E). Furthermore, the retinoic acid degrading enzyme *Cyb26b1*, which is required for the maintenance of dormant HSCs (Schönberger et al., 2022), displays elevated noise in aged LT-HSCs. Given the reduced proliferative capacity and the myeloid lineage bias of aged LT-HSCs, variability of these classes of genes could indicate the presence of differentially quiescent and lineage-biased sub-states, whereas young LT-HSCs persist in a more homogenous state.

### *Dlk1* is a marker of quiescence and enhanced self-renewal of aged HSCs

To further investigate this hypothesis, we focused on *Dlk1*, the gene with the strongest noise increase in aged versus young LT-HSCs (Figure 5D). *Dlk1* encodes a non-canonical Notch ligand which has been reported to be overexpressed in human hematopoietic CD34+ stem and progenitors from myelodysplastic syndrome patients (Sakajiri et al., 2005). In the ageing HSC dataset (Hérault et al., 2021), *Dlk1*+ and *Dlk1*-cells intermingled in the UMAP and did not give rise to separate clusters (Figure 6A). Differential gene expression analysis between *Dlk1*+ and *Dlk1*-LT-HSCs (Methods, Figure 6B) revealed only few differentially expressed genes such as the LT-HSC marker *Meg3* (Sommerkamp et al., 2019). To characterize functional differences of *Dlk1+* and *Dlk1*-LT-HSCs in more detail, we FACS-purified Dlk1+ and Dlk1-Lineage^-^ Kit^+^Sca1^+^CD150^+^CD48^-^CD34^-^ HSCs from the bone marrow of aged (18 months old) mice and performed scRNA-seq by mCEL-Seq2 (Herman et al., 2018). Gene expression analysis confirmed up-regulation of *Dlk1* mRNA in sorted Dlk1+ LT-HSCs (Figure 6C-D and S6A-B). Although clustering failed to resolve *Dlk1*+ and *Dlk1*-LT-HSCs, differential gene expression analysis between sorted Dlk1+ and Dlk1-LT-HSCs further confirmed up-regulation of *Meg3* and revealed significantly increased expression of the cell cycle inhibitor *Cdkn1a* and the Sulfotransferase 1A1 (*Sult1a1*) in Dlk1+ LT-HSCs (Figure 6E). *Sult1a1* was described as a marker of the previously identified molecular overlapping (MolO) population enriched in functional HSCs obtained by four different isolation methods (Wilson et al., 2015). These observations corroborate our sorting strategy for the two sub-populations. By FACS analysis of bone marrow cells isolated from mouse groups at different ages, we discovered that the fraction of Dlk1+ cells within the LT-HSCs compartment continuously increased with age (Figure 6F and S6C-D) and positively correlated with myeloid bias (Spearman’s *ρ*=0.80) in the bone marrow of ageing mice (Figure 6G and S6E).

**Figure 6.**
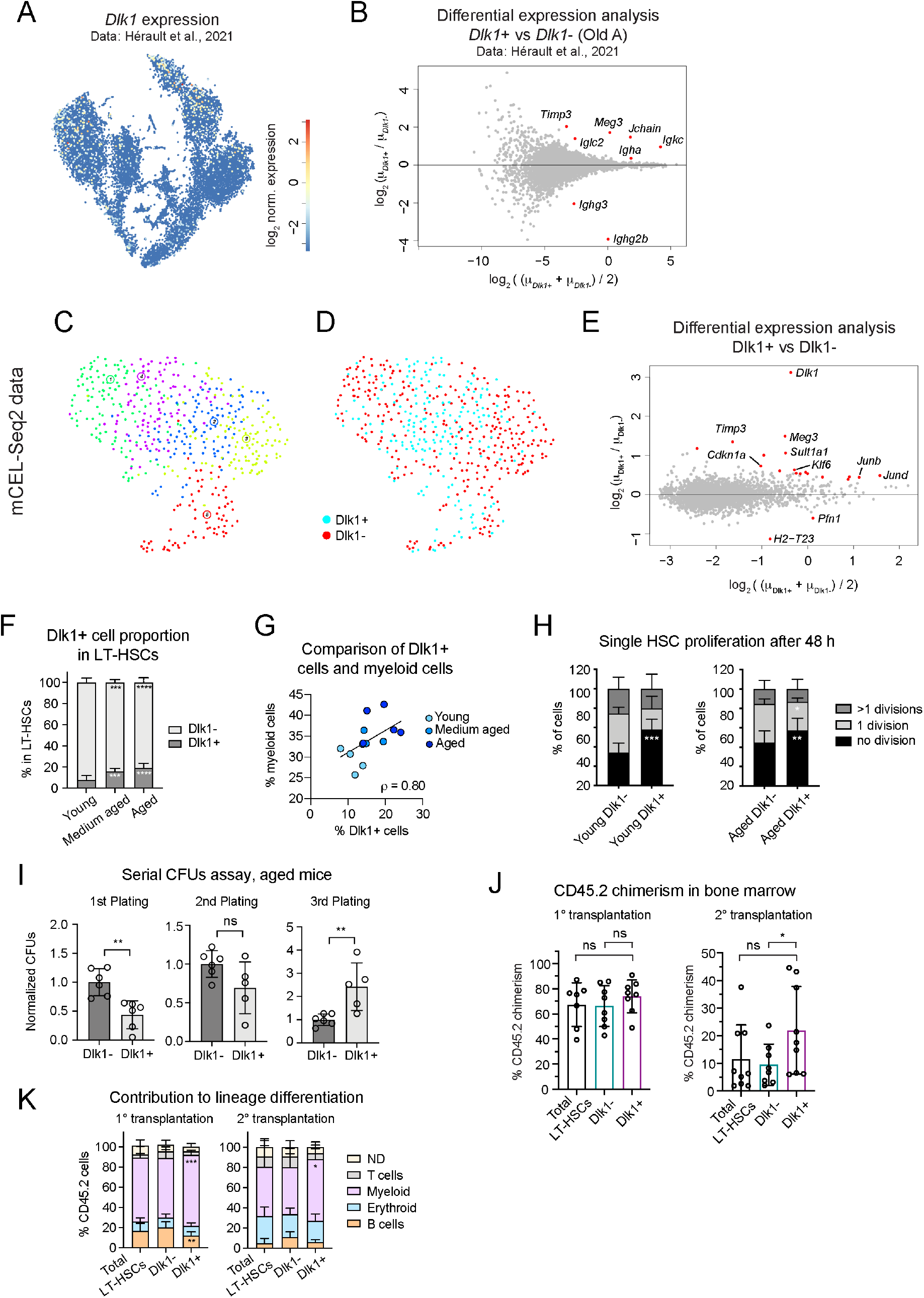
*Dlk1* is a marker of quiescence and enhanced self-renewal in aged HSCs. (A) Expression of *Dlk1* in the dataset from Hérault et al., 2021 (see Figure 5). (B) Differentially expressed genes between *Dlk1+* and *Dlk1*-cells across aged LT-HSCs, (batch A, cluster 1 in Figure 5A). Threshold values: logFC > 1.25, padj < 0.05 (C) UMAP representation of mCEL-Seq2 data of Dlk1+ and Dlk1-LT-HSC populations purified by flow cytometry. (D) Similar than (C), but highlighting Dlk1+ and Dlk1-LT-HSC sorted cells. (E) Differential expression analysis of the Dlk1+ versus Dlk1-sorted cells. Threshold values: logFC > 1.25, padj < 0.05. (F) Quantification of Dlk1+ and Dlk1-frequency among LT-HSC by flow cytometry from groups of mice with different ages (see experimental set up in Figure S6C). Error bars indicate standard deviation. (G) Comparison between the percentage of Dlk1+ cells in LT-HSCs and the percentage of myeloid cells in bone marrow, corresponding to the experiment in Figure S6C (see also Figure S6E). Spearman’s *ρ*=0.80. (H) Single cell proliferation assay showing the number of cell divisions in LT-HSCs from young (left, 3 months old) and aged (right, 17-18 months old) mice (n=3). Error bars indicate standard deviation. (I) Serial colony-forming unit assays (CFUs) with cells isolated from aged mice (17-18 months old, n=2). Error bars indicate standard deviation. (J) Percentage of CD45.2 chimerism in bone marrow 16 weeks post transplantation, showing primary (left) and secondary (right) transplantations (see experimental set up in Figure S6F). Error bars indicate standard deviation. pval: ns>0.05, *≤0.05 (one sided t-test). (K) CD42.5 lineage contribution in the bone marrow 16 weeks post transplantation, showing primary (left) and secondary (right) transplantations. Error bars indicate standard deviation. ND: non-differentiated. Statistical test in (F), (H), (I) and (K): two-way ANOVA test; pval: ns>0.05, * ≤0.05, ** ≤0.01, *** ≤0.001, **** ≤0.0001.

To better understand the specific phenotype of Dlk1+ LT-HSCs, we performed a 48h *in vitro* single-cell proliferation assay and observed delayed cell cycle entry compared to Dlk1-LT-HSCs (Figure 6H). To test *in vitro* self-renewal of the two sub-populations, we performed a serial colony-forming unit (CFU) assay. Dlk1+ LT-HSCs exhibited significantly higher CFU capacity than Dlk1-LT-HSCs after the third re-plating (Figure 6I), suggesting enhanced selfrenewal. To assess self-renewal capacity and lineage output *in vivo*, we performed serial transplantations of Dlk1+, Dlk1-, or total LT-HSCs into irradiated young recipients (Figure S6F-G, Methods). After 16 weeks of secondary transplantations Dlk1+ LT-HSC recipients exhibited significantly elevated chimerism in the bone marrow compared to Dlk1-LT-HSC recipients (Figure 6J). While we also observed elevated chimerism in the peripheral blood (Figure S6H) and an increased HSC frequency among donor-derived cells (Figure S6I) upon secondary transplantations, the difference to Dlk1-LT-HSCs recipients did not reach significance due to the large variability across individual animals. Moreover, a significantly increased myeloid lineage output was observed in the bone marrow of Dlk1+ versus Dlk1-LT-HSC recipients (Figure 6K).

Together, the *in vitro* and *in vivo* analyses indicate a more quiescent state and increased selfrenewal capacity of Dlk1+ LT-HSCs, although large variability across animals makes it difficult to quantify this effect *in vivo*. The significantly increased myeloid output in the bone marrow of Dlk1+ LT-HSC recipients is consistent with an intrinsic myeloid bias of these cells. Together with the observed correlation of Dlk1+ LT-HSC frequency and age-dependent myeloid bias (Figure 6G), we conclude that expansion of Dlk1+ LT-HSCs in the bone marrow of ageing mice contributes to the known age-related myeloid bias.

Hence, differential gene expression noise has enabled the identification of a sub-type of ageing LT-HSCs with distinct functional properties, which cannot be resolved by conventional differential gene expression analysis or clustering methods.

## Discussion

VarID2 establishes a method for the quantification of local gene expression noise in cell state space. We acknowledge that the residual variability *ε* may not be entirely free of marginal gene-specific technical noise components. However, changes in residual variability across the cell state manifold should be unaffected by such technical components, as long as noise is independent of the mean expression. In practice, the parameter *γ* driving the strength of the Cauchy prior should be adjusted such that a dependence of the average residual noise *ε* on the total UMI count per cell is eliminated.

The ability of VarID2 to quantify biological noise across neighborhoods of tens to hundreds of thousands of cells yields unprecedented insights into the dynamics of gene expression noise along differentiation trajectories of complex multilineage systems such as bone marrow hematopoiesis. This constitutes an important angle that cannot be addressed with currently available computational methods.

Consistent with a previous study measuring increased noise levels of nuclear versus cytoplasmic mRNA for ~900 genes in HeLa cells and freshly isolated primary keratinocytes (Battich et al., 2015), our study confirms a general, genome-wide increase of biological noise in the nucleus versus the cytoplasm across multiple cell types found in human peripheral blood. Therefore, as suggested previously on a limited scale (Battich et al., 2015) nuclear export is indeed likely to confer a noise buffering function on a genome-wide level with similar effect size across diverse cell types.

Making use of scRNA-seq and scATAC-seq measurements from the same cell, we identified two classes of genes with fundamentally different noise dynamics. We hypothesize that class A genes are regulated by an on/off switch lacking precise control of expression levels, whereas transcriptional levels of class B genes need to be tightly controlled. Alternatively, a correlated increase of noise and expression of class A genes could be explained by variability of extrinsic signals, e.g., related to immune cell activation, which would be consistent with the observed enrichment of immune signaling pathways among class A genes.

It requires further investigation to explain how these differential noise characteristics are regulated on the molecular level. We provide a starting point by demonstrating that different peaks within a given gene locus of class B genes are correlated with expression and noise, respectively. However, we were unable to identify global regulators of this behavior by motif analysis (data not shown).

By enabling analysis of noise dynamics during differentiation, VarID2 provides deeper insights into general properties of single-cell transcriptomes, and expands the scope of earlier work describing dynamics of transcriptome entropy during differentiation (Grün et al., 2016; Guo et al., 2016; Teschendorff and Enver, 2017). These studies consistently showed that stem cells maximize transcriptome, signaling, or pathway entropy compared to more differentiated states.

For the hematopoietic system, one of the best studied model systems for multilineage differentiation of stem cells, we reveal minimal noise levels in LT-HSCs, indicating that the quiescent state is transcriptionally homogenous.

However, we demonstrate that transcriptional noise in LT-HSCs increases with age whereby it always remains lower than in more differentiated progenitors, arguing for lower transcriptional fidelity and/or the emergence of transcriptionally similar sub-states of LT-HSCs with age, which cannot be resolved by conventional clustering approaches. Our discovery of Dlk1+ LT-HSCs, which exhibit higher self-renewal potential and myeloid bias than their Dlk1-counterpart, and which occur at increased frequency with age, provides evidence for the latter hypothesis. The correlation of Dlk1+ LT-HSC frequency with myeloid lineage frequency in the bone marrow upon ageing, in conjunction with the cell-intrinsic myeloid bias of transplanted Dlk1+ LT-HSCs, suggests this population as a determinant of age-related myeloid bias.

Due to limited transcriptional differences between Dlk1+ and Dlk1-LT-HSCs, it is impossible to distinguish these populations directly by clustering and differential gene expression analysis, highlighting that gene expression noise analysis can uncover functionally distinct sub-types in seemingly homogenous cell popualtions.

### Limitations and future directions

In order to facilitate noise inference for tens to hundreds of thousands of cell neighborhoods, VarID2 relies on maximum a posterior inference, i.e., on the inference of the most likely noise value of the posterior distribution. Hence, to account for statistical variation of this parameter, VarID2 has to rely on the distribution of MAP estimates across similar neighborhoods. We approach this here by performing a non-parametric Wilcoxon test to compare cell states, e.g., different clusters on a differentiation trajectory. Although computationally expensive, future developments could attempt full Bayesian inference of posterior parameter distributions for each neighborhood permitting to assess the uncertainty of local estimates. Another limitation is a missing link of our *ε* estimates to parameters of a mechanistic model of transcription such as the random telegraph model of transcriptional bursting (Friedman et al., 2006). Determining transcriptional parameters such as burst size and frequency relies on the validity of the underlying assumptions of the model, which could be different from gene to gene. Moreover, in the current setting we did not consider allelespecific quantification, which would require crossing of different genotypes and sufficient read coverage (Larsson et al., 2019). Nonetheless, the derivation of kinetic parameters of transcription represents an interesting future extension of VarID2.

### Conclusion

We here introduced VarID2, a novel method for the quantification of gene expression noise dynamics in cell state space, and demonstrate that noise dynamics are informative on fundamental properties and design principles of the transcriptome space. We showed that noise signals in stem cells can reveal the existence of functionally distinct sub-states, opening new avenues for investigating how functionally distinct cell states are molecularly encoded beyond differential gene expression, and for the elucidation of the role of transcriptional noise during cell fate decision in multilineage systems.

## Acknowledgments

DG was supported by the Max Planck Society, by the German Research Foundation (DFG) (322977937/GRK2344 MeInBio, SPP1937 GA 2129/2-2, GR4980/3-1, and SFB1425-Project #422681845), by the CZI Seed Networks for the Human Cell Atlas, and by the ERC (818846 — ImmuNiche — ERC-2018-COG). NC-W was supported by the Max Planck Society, ERC-Stg-2017 (VitASTEM; 759206), the DFG SFB1425 (Project #422681845), SFB992 (Project #192904750; B07), and CIBSS-EXC-2189 (Project ID 390939984). We thank the Department of Medicine II of the University Hospital Freiburg, Germany for providing human blood samples.

## Author contributions

DG conceived the study. RER and DG developed the algorithm and performed all computational analyses. JR, JSH, and GD performed experiments. NCW supervised JR. DG supervised RER, JSH, and GD. DG supervised the project. DG and RER wrote the paper and all authors edited the paper.

## Methods

### VarID2 pipeline

The original VarID method (Grün, 2020) was improved and extended to accommodate additional functionalities. In VarID2, homogeneous cellular neighborhoods were defined similarly as in VarID. In brief, a k-nearest neighbor (knn) network of cells is inferred from Pearson residuals obtained after gene-specific normalization to eliminate the dependence on total UMI counts. VarID normalization consists of a negative binomial regression of total UMI counts akin to (Hafemeister and Satija, 2019) followed by averaging regression coefficients across genes of similar expression using LOESS. In VarID2, the initial fit is performed as Poisson regression followed by a maximum likelihood inference of the dispersion parameter. As a computationally inexpensive alternative, a recently proposed analytical normalization method (Lause et al., 2021) was implemented, yielding qualitatively similar results to the full negative binomial regression. Briefly, in this normalization scheme, the regression slope coefficient equals 1 and the offset corresponds to the natural logarithm the total UMI count, if the dependent variable is the natural logarithm of a gene’s UMI count. The dispersion parameter can either be set to a fixed value, or, alternatively, inferred by maximum likelihood. After normalization, nearest neighbors are obtained by a k-d tree search based on Euclidean distance of Pearson residuals in a PCA reduced space. The number of PCs to include is inferred from an elbow criterion, requiring that the difference in explained variability upon increasing the number of PCs by one is within one standard deviation across all changes upon further increasing the number of PCs (up to a maximum of 100).

The links between a central cell and each of its *k* nearest neighbors are then tested against a negative binomial background model of UMI counts, and links to inferred outlier cells are pruned in order to obtain homogenous local cell state neighborhoods. In contrast to VarID, where the background distribution was inferred from a global mean-variance dependence of UMI counts across all genes, VarID2 constructs these background models locally, to better account for local variations in technical noise. Furthermore, to safeguard against false positive outliers due to sampling dropouts, a pseudocount of one was added to all UMI counts. The link probability is calculated as the geometric mean of the Bonferroni-corrected link probabilities of the top three genes after ranking genes by link probability in increasing order and adding a pseudocount of 10^-16^.

VarID2 also offers the possibility to estimate the a parameter, i.e., the weight of the central cell when averaging across a neighborhood for constructing the background model. The a parameter can be estimated in a self-consistent local way, requiring that the local average does not deviate more than one standard deviation from the actual expression in the central cell.

Clustering on the pruned knn network is performed by community detection. VarID2 offers to perform Leiden clustering (Traag et al., 2019) clustering with adjustable resolution parameter in addition to Louvain clustering.

In VarID2 transition probabilities between two clusters are calculated as the geometric mean of the individual link probabilities connecting the two clusters (calculated as in VarID).

Finally, apart from batch correction within the negative binomial regression framework, we integrated Harmony batch correction (Korsunsky et al., 2019). Furthermore, VarID2 facilitates pseudotime analysis along inferred lineages by integrating slingshot (Street et al., 2018).

The central noise model has been revised as outlined in the next paragraph in order to facilitate the quantification of residual biological noise.

### VarID2 noise model

In order to quantify the actual biological variability across homogenous cellular neighborhoods, we propose a statistical model that deconvolutes defined components of variability.

The UMI count *X_i,j_* detected in gene *i* and a central cell *j* within a given homogeneous neighborhood follows a negative binomial distribution, with parameters mean *μ_i,j_* and dispersion parameter *r_i,j_*:

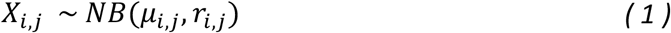

The variance of this distribution is given by

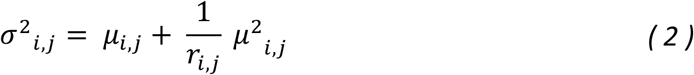

We used the definition of a Negative Binomial distribution as a Gamma-Poisson mixture. This way, the transcript counts *X_i,j_* follow a Poisson distribution with rate parameter *λ_i,j_*, which in turn follows a Gamma (*Γ*) distribution. In our model, the rate parameter *λ_i,j_* is given by

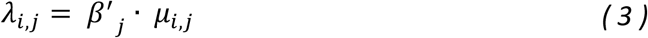

where *μ_i,j_* is the mean of the transcript counts for gene *i* across a homogenous cellular neighborhood with a central cell *j*. A homogenous cellular neighborhood *L* consist on the central cell *j* and its *k* nearest neighbors that remain after pruning: *L* = {*j, j*_1_, *j*_2_,…, *j_k_*}.

*β′_j_* is a cell specific normalization term involving the local variation in total transcript counts. Variability in total UMI counts across nearest neighbor cells are caused by technical cell-to-cell variability in sequencing efficiency and by variations in cell size or RNA content. We encompass all these sources of variability in a global term since we are interested in quantifying residual gene-specific variability.

Therefore, *β′_j_* corresponds to

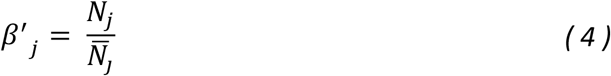

With *N_j_* representing a vector of total transcripts per cell within a neighborhood *L* with central cell *j*, and 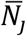 is the average of these quantities.

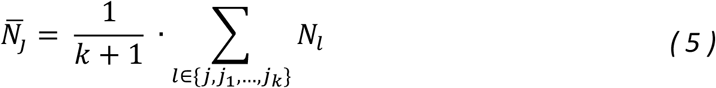

Similarly, as in (Grün et al., 2014) we propose *β′_j_* to follow a Gamma distribution with shape parameter 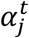 and rate parameter 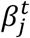. By definition it follows that

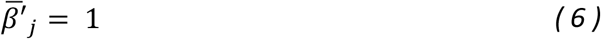

and, hence,

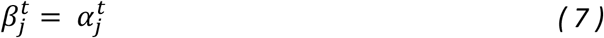

The parameter 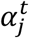 is first determined by a maximum likelihood fit of a Gamma distribution to the normalized total transcript counts, which are only marginally affected by Poissonian sampling noise due to the high magnitude of these values:

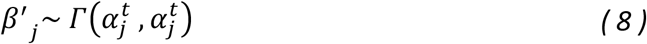

To fit a Poisson-Gamma model capturing the technical noise components defined above, and the residual biological variability, we include an inflation term *ε′_i,j_* that accounts for the biological variability of gene *i* in the neighbourhood of cell *j*.

Consequently, the rate parameter *λ_i_* follows a Gamma distribution

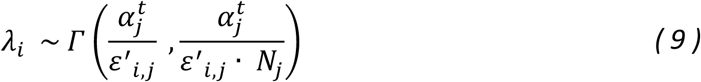

The negative binomial distribution for transcript counts *X_i,j_* is thus determined as the corresponding Poisson-Gamma mixture

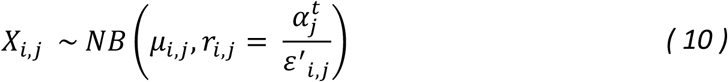

with *μ_i,j_* indicating the mean transcript counts per gene *i* across local neighbourhoods with central cell *j*.

The variance is given by

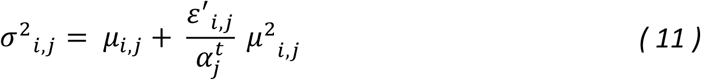

The second term in **Equation 11** can be split into the total UMI count variability contribution 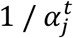 and the residual variability defined as 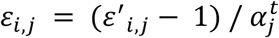, which scales from 0 to ∞. For convenience, we use 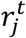 instead of 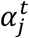, to denote the technical dispersion parameter and rewrite the variance:

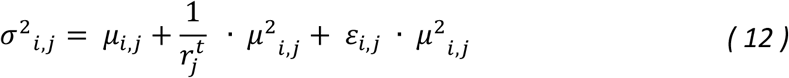

This expression encompasses the two sources of technical variability described in (Grün et al., 2014). The mean *μ_i,j_* quantifies the Poissonian noise, in which the variance scales proportionally as a function of the mean. The second technical source of variability depends on differences in sequencing efficiency or cell size and RNA content, which is captured by 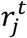. Therefore, we assume that the residual variability, denoted by *ε_i,j_*, corresponds to the biological noise.

### Implementation

Since inference of the full posterior distribution by rejection sampling across all individual neighborhoods would be computationally intense, we applied Maximum a posteriori (MAP) estimation for inferring the biological variability parameter *ε_i,j_* that maximizes the posterior distribution:

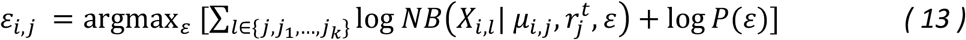

The mean expression *μ_i,j_* is calculated as the arithmetic mean of UMI counts per gene *i* across all cells *l* within local neighborhoods *L*. Alternatively, we inferred both *ε_i,j_* and *μ_i,j_* by MAP estimation, resulting in *μ_i,j_* posterior estimates highly correlated to the arithmetic mean. Therefore, we omit *μ_i,j_* from the optimization in order to reduce run time.

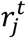 is equivalent to the shape parameter 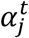 when fitting *β′_j_* (defined in **Equation 4**) by a *Γ* distribution.

We propose a Cauchy distribution as a weakly informative prior distribution *P*(*ε*) in order to regularize *ε_i,j_* posterior estimates for genes with low expression levels (Figure S1A). This prior will favour low-noise estimates in case of weak statistical support of the data, i.e., low relative likelihood.

We selected parameters of the Cauchy distribution by testing our method with a simulated dataset. In general, the location parameter *x*_0_ is set to zero and for the scale parameter we choose *γ* = 0.5 or 1.

### Determination of differentially noisy genes

To determine differentially noisy genes between two clusters, VarID2 applies a similar strategy as used in VarID. Briefly, a Wilcoxon rank-sum test of the noise levels in two clusters is performed. To mitigate the impact of the presence of only a small number of cells with non-zero noise estimates, a pseudocount sampled from a uniform distribution on [0:1] is added to each cell beforehand. Moreover, to account for reusing information across connected cells, the p-value is conservatively scaled up by the number of nearest neighbours after Bonferroni-correction across all genes and the final value is capped by 1.

### Data simulation

We generated a simulated data set with 34,390 genes and 100 cells, corresponding to a homogenous neighbourhood. Random transcript counts were sampled from a negative binomial distribution with mean *μ_i_* and dispersion parameter *r_i_* based on a reference dataset (Tusi et al., 2018). The mean *μ_i_* was defined as the average of transcript counts per gene across the reference dataset and multiplied by the parameter *α_j_*, a cell-specific term accounting for the discrepancies in sequencing efficiency across individual cells. *α_j_* was generated by sampling random values from a *Γ* distribution with both shape and rate parameters equal to 2.

We estimated the dispersion parameter per gene from the reference dataset as: 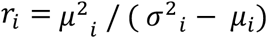. Using this set of *r_i_* values, we took the 0.2, 0.4 and 0.8 quantiles to define three levels of noise: high, medium and low, respectively. This way, we generated a simulated dataset whose genes have three levels of variability and their expression cover a broad range (See Figure S1A).

### Gene expression variability estimation with BASiCS

We applied BASiCS (Eling et al., 2018; Vallejos et al., 2015) to the simulated dataset by using the implementation with regression model and without spike-in, and choosing default parameters.

### Analysis of publicly available datasets

For analysis of public datasets, feature per barcode matrices with raw counts were retrieved from NCBI Gene Expression Omnibus (GEO; https://www.ncbi.nlm.nih.gov/geo/) or from the 10x Genomics website (https://support.10xgenomics.com/single-cell-multiome-atac-gex/datasets). See details in Supplementary Table S1. We ran the VarID2 pipeline implemented in the RaceID3 package (v0.2.5) for the analysis of single-cell transcriptome data. Unless otherwise indicated, we processed the data as follows: cells with less than 1000 UMI counts were filtered out. Genes that do not have at least 5 UMI counts in at least 5 cells were discarded. Mitochondrial genes, ribosomal genes, predicted genes with Gm-identifier and genes correlated to these classes were removed (CGenes argument in filterdata function). The pruned k-nearest neighbor (knn) network of cells was computed with the pruneKnn function, with number of neighbors set to 25. Clustering was performed with the Leiden algorithm for community detection (Traag et al., 2019) implemented in the graphCluster function. t-SNE or UMAP dimensional reduction representations were computed with comptsne and compumap functions with default parameters.

For local noise quantification, we adjusted the value of the prior parameter *γ* based on several criteria: closeness to the simulated ground truth and reduced standard deviation of the noise estimates (See Figure S1C). We selected low values of *γ* (around 0.5 and 1) in order to avoid inflation of noise estimates for lowly expressed genes. We also assessed the absence of correlation between total UMI count and noise estimates (See Figure S4C). Cellular noise was defined as the mean noise estimates per cell, averaging across all genes.

#### Human PBMCs, Multiome Assay and scRNA-seq Assay

##### Transcriptomics data

snRNA-seq and scRNA-seq datasets were individually analyzed with the VarID2 pipeline. To be consistent with Seurat pipelines, cells with less than 1000 or more than 25,000 UMI counts were discarded. Only mitochondrial genes, ribosomal genes and predicted genes with Gm-identifier were filtered (FGenes argument in filterdata function). To keep a comparable number of clusters between both datasets, the Leiden resolution was adjusted to 2 (nuclei data) and 1.5 (cell data). For noise estimation, we set the prior parameter *γ* = 1.

##### Batch correction with Harmony and comparison of gene expression noise between datasets

Matrices with raw UMI counts of snRNA-seq and scRNA-seq datasets were pooled together. Only the cells passing quality filters after VarID2 analysis were used. Batch correction with Harmony (Korsunsky et al., 2019) was performed with the implemented function in the Seurat package (Hao et al., 2021; Stuart et al., 2019), by using default parameters and following the vignette: https://portals.broadinstitute.org/harmony/SeuratV3.html. The resulting cluster labels were used to compare cell populations, according to the following annotation: 0: CD4 memory T cells; 1, 3, 11: CD14 monocytes; 2: CD4 naïve T cells; 4: CD8 naïve T cells; 5, 7: B cells; 6: CD8 effector memory T cells 1 (TEM1); 8: natural killer cells (NK); 9: CD8 effector memory T cells 2 (TEM2); 10: CD16 monocytes.

##### Noise quantification for smFISH data and comparison with sn- and scRNA-seq data

In order to quantify noise from smFISH data, we assumed that the counts follow a negative binomial distribution with variance

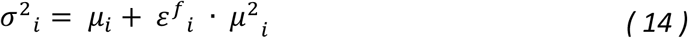

where *ε^f^_i_*, is the reciprocal of the dispersion parameter and analogous to the noise parameter *ε_i,j_* obtained from our VarID2 method. Unlike scRNA-seq, smFISH is only marginally affected by technical variability of signal detection across cells. Furthermore, since we applied smFISH to CD8 T cells, we do not expect substantial variation in cell size and RNA content. Therefore, we omitted the dispersion parameter associated with total UMI count variability, which cannot be inferred based on the quantification of individual genes.

We computed the ratio of noise estimates between nuclear and cellular compartments and estimated the error based on the standard error of the mean.

##### Analysis of scATAC-seq data

scATAC-seq data was analyzed with Signac package (v1.2.1) and Seurat (v4.0.3) (Hao et al., 2021; Stuart et al., 2019). We followed the WNN vignette of 10x Multiome, RNA + ATAC to facilitate the joint analysis of both modalities. Gene activities, defined as the sum of detected fragments across all peaks located in the gene body and 2 kb upstream of the transcriptional start site, were computed with default parameters with GeneActivity function. Peak to gene links (Ma et al., 2020) were computed with LinkPeaks function with default parameters. For expression – peak links, the Assay with expression data was used. For noise – peak links an additional Assay denoted “Noise” was created within the Seurat object.

Differential accessibility tests were performed with FindMarkers function with the logistic regression method (Ntranos et al., 2019). In brief, this method establishes a logistic model based on fragment abundance of a given feature and performs a likelihood ratio test by comparing this to a null model.

#### Murine hematopoietic progenitor cells (Dahlin et al.,2018)

WT and *Kit* mutant W^41^/W^41^ samples were analyzed individually. Low quality cells with less than 2000 UMI counts were removed. We used 50 nearest-neighbors for inference of the pruned knn network. Leiden resolution: 1.5. Prior parameter *γ* = 0.5.

#### Hematopoietic progenitors from young and aged mice (Hérault et al., 2021)

Louvain clustering was performed. t-SNE perplexity = 200, Prior parameter *γ* = 0.5.

##### Transition probabilities

Transition probabilities were estimated by VarID2 (Grün, 2020). Based on the connections in the pruned knn, probabilities of individual links connecting cells from two different clusters are estimated. Transition probabilities between two clusters correspond to the geometric mean of the individual link probabilities connecting the two clusters.

##### Quadratic programming to identify similarities between two datasets

We employed quadratic programming for mapping corresponding cell populations between WT and W^41^/W^41^ samples of the HSC data (Dahlin et al., 2018). We represented the cluster medoids of one dataset as a linear combination of the medoids from the other dataset. Subsequently, we optimized the weights for all cluster medoids under the constraints that they are greater or equal than zero and that they sum up to one. We solve this optimization problem with the QP function of the quadprog R package.

##### Differential Expression Analysis

Differential expression analysis was computed with the diffexpnb function of the RaceID3 (v0.2.5) algorithm. Detection of differentially expressed genes between specific groups of cells was performed with a similar method as previously reported (Anders and Huber, 2010). In brief, a negative binomial distribution which captures the gene expression variability for each group of cells if inferred based on a background model of the expected transcript count variability estimated by RaceID3 (Herman et al., 2018). Based on the inferred distributions, a *P* value for the significance of the transcript counts between the two groups of cells is estimated and multiple testing corrected by Benjamini-Hochberg method.

##### Pathway Enrichment Analysis

Pathway enrichment analysis was performed with the enrichPathway function from the ReactomePA R package (Yu and He, 2016) or compareCluster from clusterProfiler R package (Yu et al., 2012), with p-value cut off = 0.05 and multiple testing corrected by the Benjamini-Hochberg method. Input were ENTREZ gene IDs of genes selected by differential expression analysis or detection of differentially noisy genes.

##### Data availability

mCELseq2 data were deposited on the Gene Expression Omnibus database, with accession number GSE185637.

##### Code Avalability

VarID2 is part of the RaceID R package (v0.2.5), available on github https://github.com/dgrun/RaceID3_StemID2_package and on CRAN.

### Experimental Section

#### Experimental models and subject details

##### Mouse

Experiments were performed with wild-type C57BL/6J male mice, obtained from in-house breedings or ordered from JAX. Mice were maintained under specific-pathogen-free conditions within the animal facility of the Max Planck Institute of Immunobiology and Epigenetics. Protocols for animal experiments were approved by the review committee of the Max Planck Institute of Immunobiology and Epigenetics and the Regierungspräsidium Freiburg, Germany.

##### Human blood samples

Blood samples were obtained from healthy donors recruited at the Department of Medicine II of the University Hospital Freiburg, Germany.

Written informed consent was given by all donors prior to blood donation. Peripheral blood mononuclear cells (PBMCs) from EDTA-anticoagulated participant blood were isolated by density gradient centrifugation using Pancoll (Pan-Biotech).

##### Cell suspensions and flow cytometry

Murine bone marrow (BM) cells were isolated from pooled femura, tibiae, hips, ilia, and vertebrae by gentle crushing in PBS using a mortar and pistil. Erythrocyte lysis was performed using ACK Lysing Buffer. To enrich for lineage negative (Lin^-^) cells Dynabeads Untouched Mouse CD4 Cells kit (Invitrogen) were used according to the manufacturer’s instructions. Briefly, the erylsed BM was stained for 40 min with the provided Lineage Cocktail. Labelled cells were incubated for 15 min with polyclonal sheep anti-rat IgG coated Dynabeads (provided in the kit). Subsequently, labelled Lin+ cells were magnetically depleted. To achieve further purification, HSCs were FACS sorted. Therefore, the depleted cell fraction was stained for 30 min to 1 h using the following monoclonal antibodies: anti-lineage [anti-CD4 (clone GK1.5), anti-CD8a (53-6.7), anti-CD11b (M1/70), anti-B220 (RA3-6B2), anti-GR1 (RB6-8C5) and anti-TER119 (Ter-119)] all PE-Cγ7; anti-CD117/c-Kit (2B8) in BV711; anti-Ly6a/Sca-1 (D7)-APCCy7; anti-CD34 (RAM34) in AF700; anti-CD150 (TC15-12F12.2) in PE/Dazzle; anti-CD48 (HM48-1) in BV421. Monoclonal antibodies were purchased from eBioscience, BioLegend or MBL. Either DLK1+ or DLK1-HSCs were sorted. Cells were sorted into Stem Pro®-34 SFM (Life Technologies) for further experiments.

##### Single cell (SiC) division assay

Single DLK1+ or DLK1-HSCs (Lineage^-^Kit^+^Sca1^+^CD150^+^CD48^-^CD34^-^) were FACS sorted into 72-well Terasaki plates and cultured in StemPro-34 SFM containing 50 ng/ml SCF, 25 ng/ml TPO, 30 ng/ml Flt3-Ligand, 100 ml/ml Penicillin/Streptomycin, 2 mM L-Glutamine. After 48h each well was checked manually for the number of cell divisions under the microscope: 1 cell = no division, 2 cells = 1 division, >2 cells= >1 division.

##### Serial Colony-Forming-Unit assays (CFU)s

200-400 DLK1+ and DLK1-HSCs (Lineage^-^Kit^+^Sca1^+^CD150^+^CD48^-^CD34^-^) were FACS sorted into MethoCult M3434, plated and cultured. Approximately 7 days after the first plating, number of colonies were counted and 10,000 cells were re-plated. 2nd and 3rd platings were performed 3 and 5 days, respectively, after the first re-plating. Colonies were also quantified at these time points.

##### HSC transplantation assay

600 Dlk1+, Dlk1-, or total HSCs (Lineage^-^Kit^+^Sca1^+^CD150^+^CD48^-^CD34^-^) isolated from 13-15 month old CD45.2 C57BL/6 mice were transplanted into lethally irradiated (4.5 Gray + 5 Gray) CD45.1 (Ly5.1) mice together with 5×10^5^ supportive spleen cells from 8-12 week old CD45.1/2 mice within 24h after irradiation by intravenous tail vein injection. Contribution of donor cells (CD45.2) was monitored in peripheral blood at 4, 8, 12 and 16 weeks post-transplantation. For endpoint analysis, bone marrow was analyzed at 16 weeks post transplantation to quantifiy CD45.2 chimerism and lineage contribution. For secondary transplantations, 3 × 10^6^ cells of whole bone marrow was isolated and retransplanted 16 weeks post-transplantation. CD45.2 chimerism and lineage contribution in bone marrow and peripheral blood were quantified by flow cytometry using the following antibodies: anti-CD45.1 (A20)–FITC, anti-CD45.2 (104)–PB, anti-CD11b (M1/70)-APCCy7, anti-GR1 (RB6.8C5)-APC, anti-CD8a (53.6.7)-PECy5, anti-CD4 (GK1.5)-PECy5, anti-B220 (RA3.6B2)-AF700.

##### Amplified RNA preparation from single cells using mCEL-Seq2

The CEL-Seq2 protocol with reduced volumes was used as previously described (Herman et al., 2018) and modified using the following reagents.

Instead of 1.2 μl Vapour-Lock as hydrophobic encapsulation barrier mineral oil (Sigma, M8410-100ML) was used. For cDNA first-strand synthesis, Protoscript II and Protoscript II Reaction Buffer (NEB, M0368L) as well as murine RNase-Inhibitor (NEB, M0314S) was used instead of SuperScript II reverse transcriptase, first-strand synthesis buffer and RnaseOUT. *Escherichia coli* DNA polymerase I, *E. coli* DNA ligase, RNase H (Invitrogen; 18021071) and 5 x second-strand buffer were replaced with *E. coli* DNA polymerase (NEB, M0209L), *E. coli* DNA ligase (NEB, M0205L), RNaseH (NEB, M0297S) and 10x Second Strand Buffer (NEB, B6117S) respectively.

The water volume was adjusted to adequately dilute the 10x second strand buffer. After second strand synthesis 96 wells were pooled, which results in 96 single cells per library.

The library preparation was performed as previously described (Herman et al., 2018), but by using Protoscript II, Protoscript II Reaction Buffer and murine RNase-Inhibitor as mentioned above instead of SuperScript II reverse transcriptase, first-strand synthesis buffer and RnaseOUT.

##### Quantification of transcript abundance

Paired-end reads were aligned to the transcriptome using BWA (version 0.6.2-r126) with default parameters (Li and Durbin, 2010). The transcriptome contained all gene models based on the mouse ENCODE VM9 release downloaded from the UCSC genome browser comprising 57,207 isoforms with 57,114 isoforms mapping to fully annotated chromosomes (1–19, X, Y, M). All isoforms of the same gene were merged to a single gene locus, and gene loci were merged to larger gene groups, if loci overlapped by >75%. This procedure resulted in 34,111 gene groups. The right mate of each read pair was mapped to the ensemble of all gene groups in the sense direction. Reads mapping to multiple loci were discarded. The left mate contained the barcode information: the first six bases corresponding to the cell-specific barcode, followed by six bases representing the UMI. The remainder of the left read contained a poly(T) stretch and adjacent gene sequence. The left read was not used for quantification. For each cell barcode and gene locus, the number of UMIs was aggregated and, on the basis of binomial statistics, converted into transcript counts (Grün et al., 2014).

##### smRNA FISH

Singly labelled oligonucleotides (Quasar 570 or Quasar 670) targeting *PPP1R2* and *PDCD4* mRNAs were designed with the Stellaris RNA FISH probe designer (LGC Biosearch Technologies, version 4.2) and produced by LGC Biosearch Technologies.

Naïve CD8+ T cells were isolated from the peripheral blood of two healthy donors using the Naive CD8+ T Cell Isolation kit (Miltenyi Biotec, 130-093-244) according to the manufacturer’s instructions. SmRNA FISH procedure was performed in suspension, with brief centrifugations between steps (5 min at 400 x g) to remove the supernatant. Briefly, naïve CD8+ T cells were washed once with PBS, fixed in 3.7% formaldehyde in PBS for 10 min at room temperature, and washed again twice with PBS. Cell pellet was resuspended in 200 μl of 70% ethanol, incubated at 4°C for 1 h and then washed with 200 μl of wash buffer A (LGC Biosearch Technologies, SMF-WA1-60) supplemented with 10% deionized formamide (Thermo Fisher Scientific, 4440753) at room temperature for 5 min. Cells were hybridized with 80 μl of hybridization buffer (LGC Biosearch Technologies, SMF-HB1-10) supplemented with 10% deionized formamide containing the FISH probes at a 1:100 dilution at 37°C overnight. The next day, cells were washed with 200 μl of wash buffer A supplemented with 10% deionized formamide at 37°C for 30 min and stained with wash buffer A supplemented with 10% deionized formamide and 10 μg/ml Hoechst 33342 (Thermo Fisher Scientific, H3570) at 37°C for 30 min. Cells were rinsed once with 200 μl of 2x SSC, equilibrated 5 min in base glucose buffer (2X SSC, 0.4% glucose solution, 20 mM Tris pH 8.0 in RNase-free H2O), and then incubated 5 min in base glucose buffer supplemented with a 1:100 dilution of glucose oxidase (stock 3.7 mg/ml) and catalase (stock 4 mg/ml). Cell pellet was resuspended in 10 μl of ProLong Glass Antifade Mountant (Thermo Fisher Scientific, P36984) and mounted on a glass slide with a glass coverslip.

##### Microscopy and image analysis

Z-stacks with 250-350 nm z-steps were acquired with the Cell Observer spinning disk confocal microscope from Zeiss with a 100×/1.40-numerical aperture oil objective lens and the PrimeBSI camera from Photometrics. Cells were segmented using Imaris image analysis software (Bitplane) and FISH spots within nucleus and cytoplasm were quantified.

## Supplemental Figures

**Figure S1.**
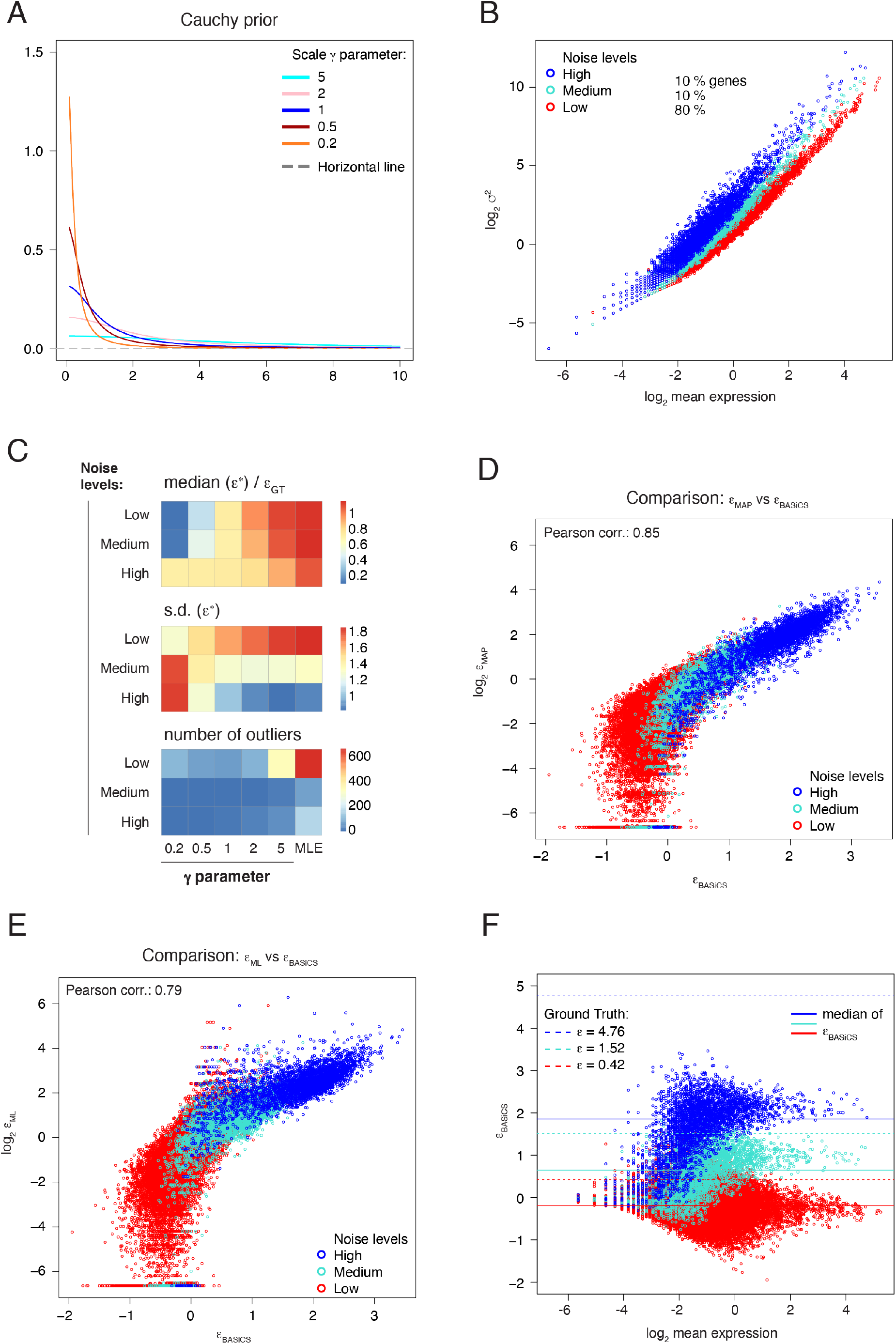
Local decomposition of gene expression noise in cell state space, Related to Figure 1. (A) Probability density function of the Cauchy distribution with different values of the scale parameter *γ*. Only positive values are considered for the Cauchy distribution. Location parameter was set to zero. (B) Variance as a function of the mean on a logarithmic scale for a simulated dataset with genes grouped into three levels of biological noise (Methods). (C) Tests for hyperparameter *γ* selection, based on the ratio: *median* (*ε*^*^)/*ε_GT_* (GT: Ground Truth); the standard deviation *s. d*. (*ε*^*^); and number of outliers, given by *ε* > *ε_GT_* + *s. d*. (*ε*^*^). *ε*^*^ estimates correspond to genes whose mean expression meets the condition: *log*_2_ (*μ_i_*) > 1. (D) Comparison of *ε_MAP_* estimates, with *ε_BASiCS_*, the dispersion parameter *ε* computed by BASiCS (Eling et al., 2018). (E) Similar to (D), but comparison of maximum likelihood estimates *ε_MLE_* and *ε_BASiCS_* estimates. (F) Scatterplot of *ε_BASiCS_* estimates as a function of the mean of the simulated dataset.

**Figure S2.**
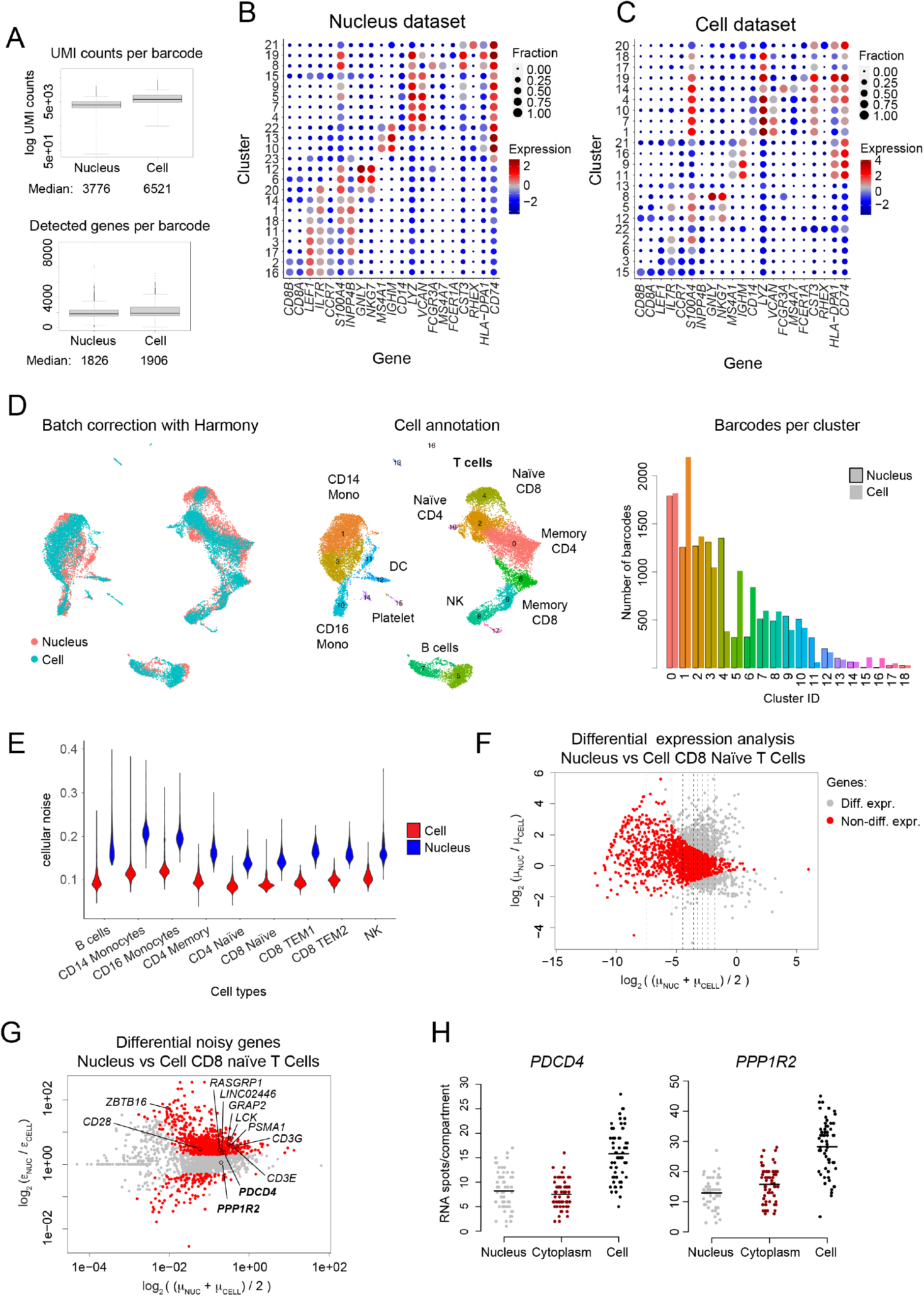
Elevated noise levels of nuclear versus whole-cell transcriptomes in human PBMCs, Related to Figure 2. (A) Number of UMI counts (top) and number of detected genes per barcode (bottom) for snRNA-seq (Nucleus) and scRNA-seq (Cell) data of human PBMCs. Boxes indicate interquartile range (IQR), and whiskers correspond to ±1.5*IQR of the box limits. Outliers beyond the whisker limits are depicted. (B) Expression of relevant marker genes across clusters in the snRNA-seq dataset. Dot size indicates the fraction of cells with positive expression and dot color highlights logarithmic expression (log2) calculated across clusters. (C) Same as (B), but expression detected in the scRNA-seq dataset. (D) Batch effect correction with Harmony. UMAP representations showing the distribution of barcodes per sample (left), cell type annotations (middles), as well as the number of barcodes per dataset assigned to each cluster (right). (E) Comparison of cellular noise across the main cell populations detected in (D). See also Figure 2E. For better visualization, outliers >0.4 are not included. (F) Differential expression analysis of CD8 naïve T cells (cluster 4 in (D)), comparing scRNA-seq versus scRNA-seq samples. Genes were split into ten equally populated bins, based on their mean expression (vertical lines) and genes with no differential expression (red dots) were selected to compare noise levels per gene (see Figure 2F). Threshold values: FC > 1.25, padj < 0.001. (G) Test of differential noisy genes in CD8 naïve T cells, comparing nuclei versus cell samples. Threshold values: fold-change (FC) > 2, adjusted *P* (padj) < 0.001. (H) Quantification of RNA spots for each cellular compartment from smFISH experiments. See also Figure 2G – H. Horizontal lines indicate the mean. DC, dendritic cells; NK, natural killer cells, TEM, effector memory T cells; Mono, monocytes.

**Figure S3.**
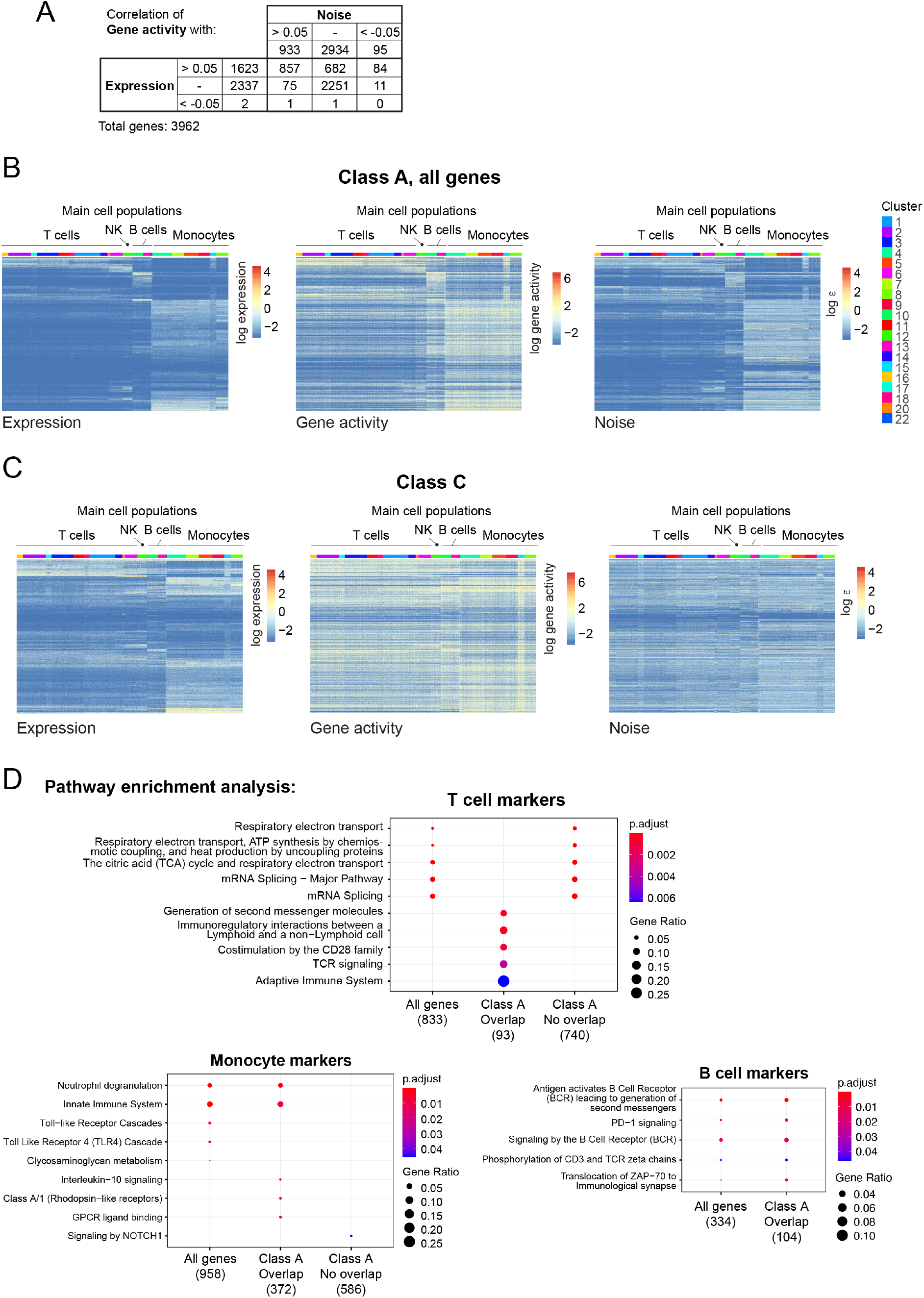
Joint analysis of chromatin accessibility and gene expression noise reveals gene modules with distinct noise regulation, Related to Figure 3. (A) Pearson correlations between gene activity and gene expression, and between gene activity and noise were computed. The contingency table shows the number of genes identified with a positive (> 0.05), negative (< −0.05) or undefined (> −0.05 & < 0.05) correlation. (B) Heatmaps showing patterns of expression, gene activity and noise of all genes belonging to class A. See also Figure 3B. (C) Similar to (B), but patterns of expression, gene activity and noise for class C genes. (D) Pathway enrichment analysis performed for marker genes of T cells, monocytes and B cells. Maker genes were defined as cell type-enriched genes by performing pairwise differential expression analyses. For each major cell population, marker genes were analyzed in three groups: all genes, marker genes belonging to Class A (“Overlap”), and marker genes not belonging to Class A (“No overlap”). For B cell marker genes, the non-overlapping group was not significantly enriched in any pathway.

**Figure S4.**
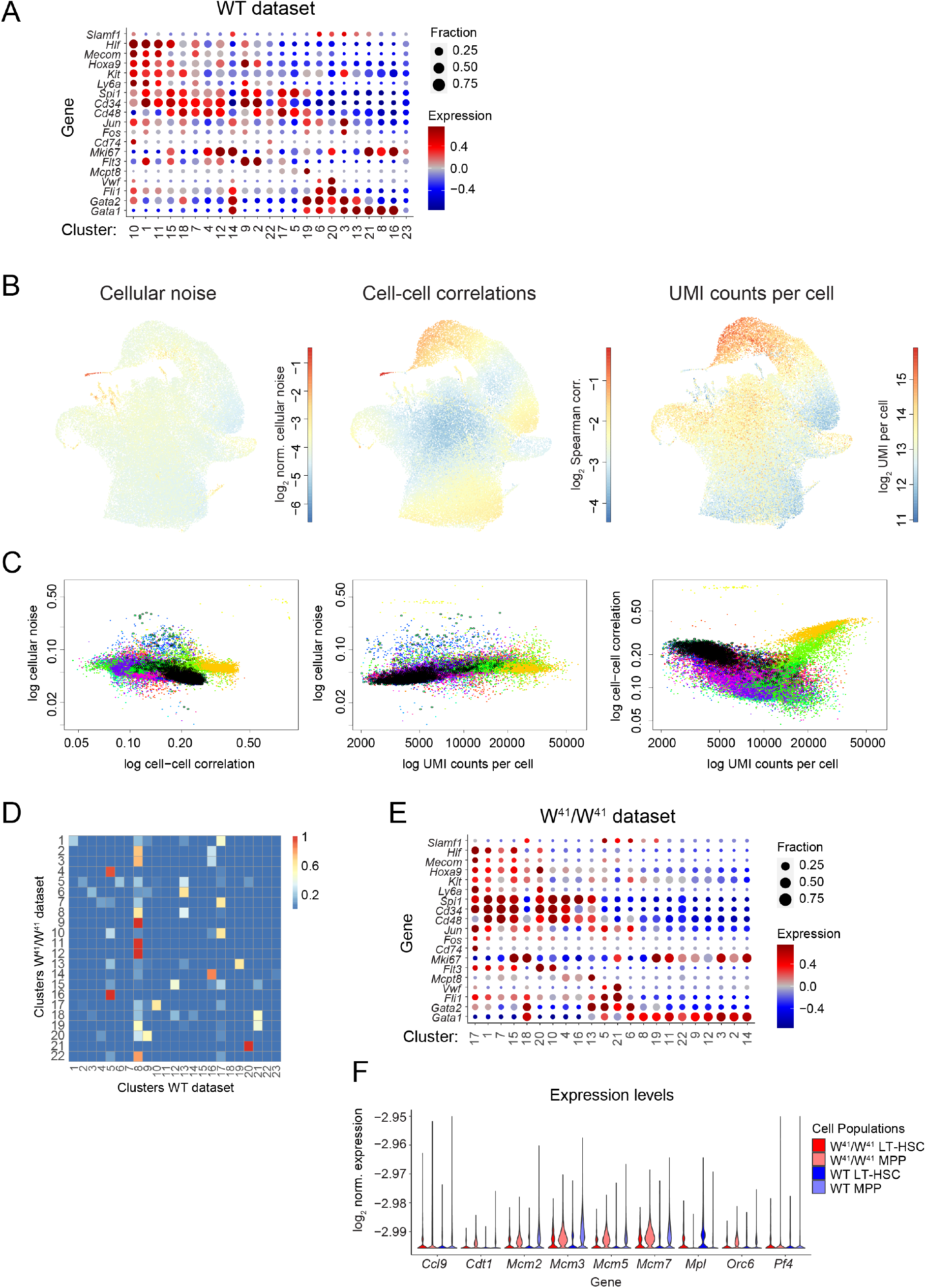
Gene expression noise increases during hematopoietic differentiation, Related to Figure 4. (A) Expression of relevant marker genes across cell clusters in the WT dataset of hematopoietic progenitors. Dot size indicates the fraction of cells with positive expression and dot color highlights expression z-score calculated across clusters. Values higher than 0.75 and lower than −0.75 are replaced by 0.75 and −0.75, respectively. See clustering in Figure 4A. (B) UMAPs highlighting cellular noise (left), local cell-cell correlations (center) and UMI counts per cells (right) across the WT dataset. (C) Scatterplots of the quantities displayed in (B). Colors correspond to the clusters in Figure 4A. Data points of the LT-HSCs (cluster 10) are highlighted with a black outline. (D) Heatmap depicting similarity weights of clusters in the W^41^/W^41^ dataset to WT cluster inferred by quadratic programming (Methods). (E) Similar as (A), but for the W^41^/W^41^ dataset. (F) Violin plot showing normalized expression of genes involved in DNA replication, similar to the genes shown in Figure 4G. Samples are separated into LT-HSC and the remaining cells, denoted MPP. For better visualization, outliers >-2.95 are not included.

**Figure S5.**
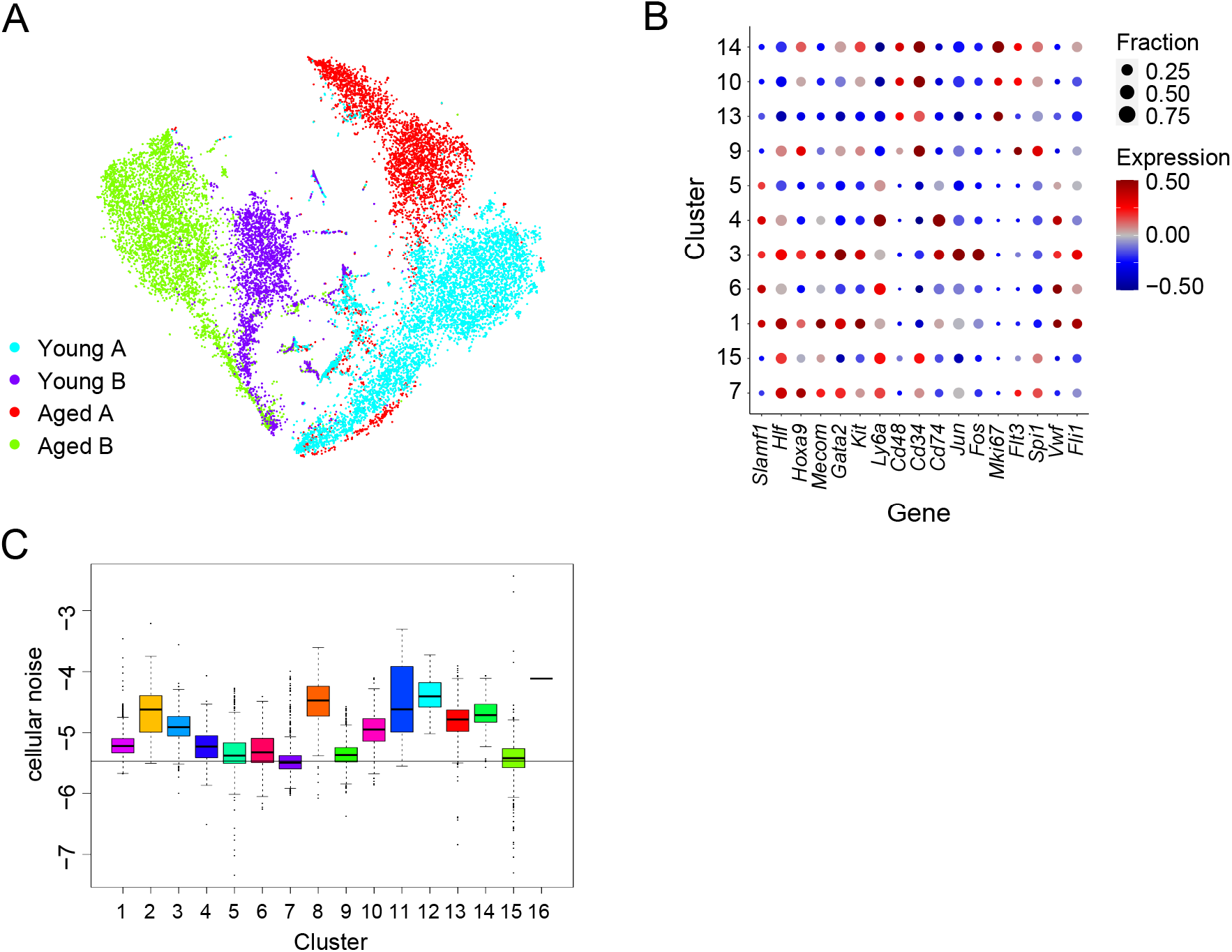
Gene expression noise increases in LT-HSCs upon ageing, Related to Figure 5. (A) t-SNE map highlighting the four samples and processed in to batches from the Hérault et al. (2021) dataset. (B) Expression of relevant marker genes across cell clusters in the dataset. See clustering in Figure 5A. Dot size indicates the fraction of cells with positive expression and dot color highlights expression z-score calculated across clusters. Values higher than 0.5 and lower than −0.5 are replaced by 0.5 and −0.5, respectively. (C) Quantification of cellular noise across all clusters in the dataset. Horizontal line corresponds to the median noise level of the LT-HSC young A population (cluster 7). Boxes indicate inter-quartile range, and whiskers correspond to ±1.5*IQR of the box limits. Outliers beyond the whisker limits are depicted.

**Figure S6.**
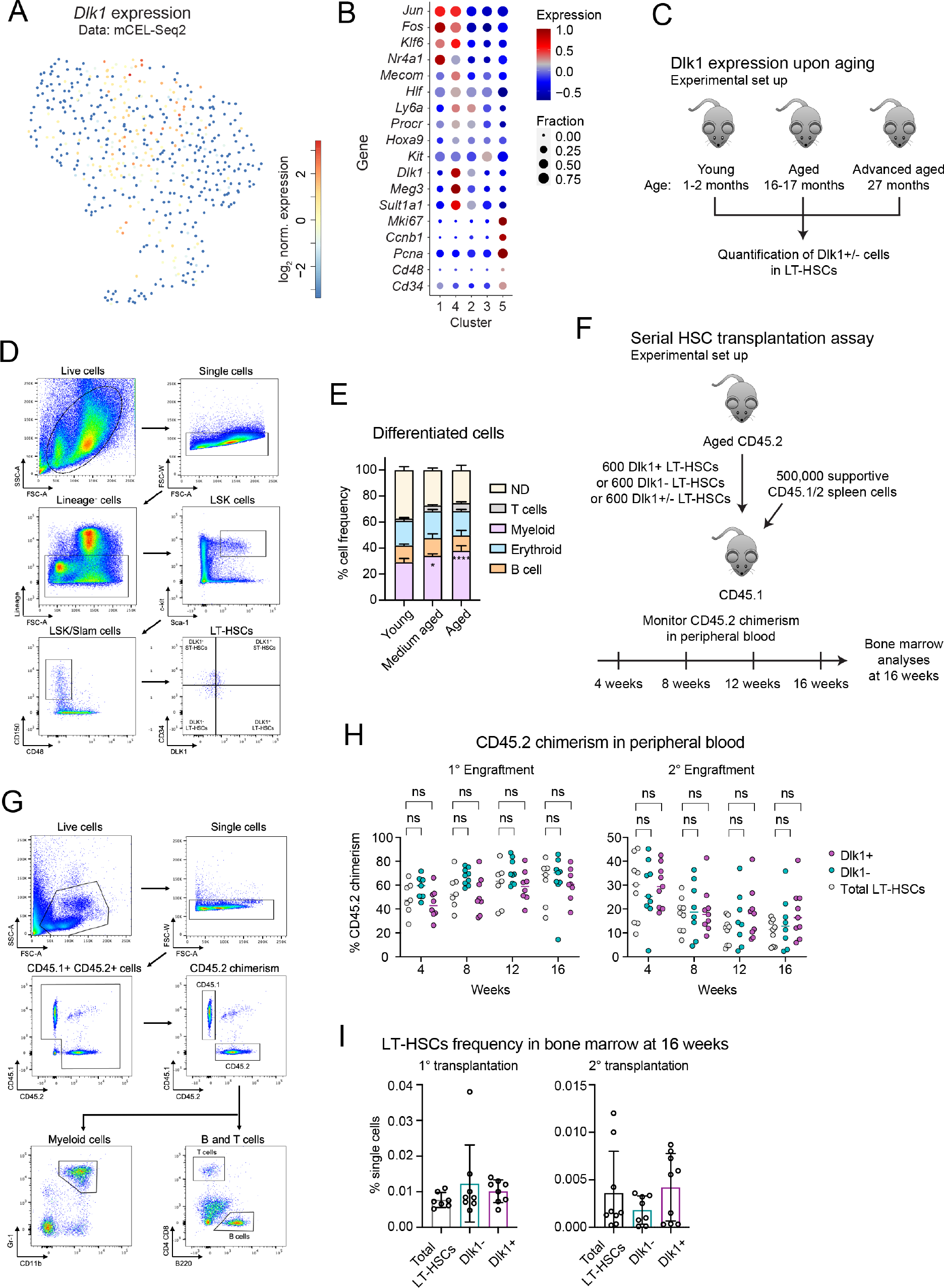
Dlk1 is a marker of quiescence and enhanced self-renewal of aged HSCs, Related to Figure 6. (A) UMAP representation highlighting expression of *Dlk1* in the mCEL-Seq2 dataset shown in Figure 6C-D. (B) Expression of relevant marker genes across cell clusters in the mCEL-Seq2 dataset. Dot size indicates the fraction of cells with positive expression and dot color highlights expression z-score calculated across clusters. Values higher than 1 are replaced by 1. (C) Experimental design for quantification of Dlk1+/- LT-HSC frequency upon aging. (D) Representative gating scheme for sorting Dlk1+/- LT-HSCs, corresponding to the experiment described in (C), see also Methods. (E) Hematopoietic lineage analysis of bone marrow for the age groups described in (C). ND: non-differentiated. Error bars indicate standard deviation. (F) Experimental design of serial HSC transplantation assay, see also Methods. (G) Representative gating scheme for monitoring the CD45.2 chimerism, corresponding to the experiment described in (F), see also Methods. (H) Percentage of CD45.2 chimerism in peripheral blood across the indicated time points for primary (left) and secondary (right) transplantations, corresponding to the experiment described in (F), see also Methods. (I) Quantification of LT-HSCs in bone marrow 16 weeks post transplantation, showing primary (left) and secondary (right) transplantations. Error bars indicate standard deviation. Barplots and scatterplots: pval: ns>0.05, * ≤0.05, ** ≤0.01, *** ≤0.001, **** ≤0.0001 (twoway ANOVA test).

**Supplementary Table S1.** List of public datasets analyzed.

| Dataset | Reference | Repository / Accession number |
| --- | --- | --- |
| Murine Kit+ hematopoietic progenitor cells from bone marrow | Tusi et al., 2018 | GEO: GSE89754 |
| Human PBMC Single Cell Gene Expression Assay (v3 chemistry) | - | 10x Genomics; https://support.10xgenomics.com/single-cell-gene-expression/datasets/3.0.0/pbmc_10k_v3 |
| Human PBMC, Single Cell Multiome ATAC + Gene Exp. Assay | - | 10x Genomics, https://support.10xgenomics.com/single-cell-multiome-atac-gex/datasets/1.0.0/pbmc_granulocyte_sorted_10k |
| Murine hematopoietic progenitor cells from bone marrow | Dahlin et al., 2018 | GEO: GSE107727 |
| Aged and young hematopoietic stem cells from murine bone marrow | Hérault et al., 2021 | GEO: GSE147729 |

